# Abundance measurements reveal the balance between lysis and lysogeny in the human gut microbiome

**DOI:** 10.1101/2024.09.27.614587

**Authors:** Jamie A. Lopez, Saria McKeithen-Mead, Handuo Shi, Taylor H. Nguyen, Kerwyn Casey Huang, Benjamin H. Good

## Abstract

The human gut contains diverse communities of bacteriophage, whose interactions with the broader microbiome and potential roles in human health are only beginning to be uncovered. Here, we combine multiple types of data to quantitatively estimate gut phage population dynamics and lifestyle characteristics in human subjects. Unifying results from previous studies, we show that an average human gut contains a low ratio of phage particles to bacterial cells (∼1:100), but a much larger ratio of phage genomes to bacterial genomes (∼4:1), implying that most gut phage are effectively temperate (e.g., integrated prophage, phage-plasmids, etc.). By integrating imaging and sequencing data with a generalized model of temperate phage dynamics, we estimate that phage induction and lysis occurs at a low average rate (∼0.001-0.01 per bacterium per day), imposing only a modest fitness burden on their bacterial hosts. Consistent with these estimates, we find that the phage composition of a diverse synthetic community in gnotobiotic mice can be quantitatively predicted from bacterial abundances alone, while still exhibiting phage diversity comparable to native human microbiomes. These results provide a foundation for interpreting existing and future studies on links between the gut virome and human health.

## Introduction

The human gut harbors a complex community of bacteria, viruses, and microbial eukaryotes that plays important roles in human health (1–3). Previous studies have largely focused on the bacterial portion of this community, but in recent years the bacteriophage (“phage”) that infect these bacteria have started to draw more attention. Advances in DNA sequencing and anaerobic culturing have led to extensive databases of gut phage genomes (4,5), as well as increasing numbers of phage isolates that can be propagated in the lab for mechanistic investigation (6,7). Phage can influence the microbiome in multiple ways. They can directly kill their bacterial hosts through lytic infection (7,8) or by inducing lysis from a temperate state (8,9). Temperate phage can also serve as important vectors of horizontal gene transfer (10), carrying cargo genes that enhance the metabolic or defense capabilities of their bacterial hosts (11). These interactions with gut bacterial ecology and evolution have been hypothesized to impact human health. Cohort studies have revealed numerous associations between the composition of the gut virome and various health-related states, including cancer treatment efficacy (2) and lifespan (12). Transplants of sterile phage-containing fecal filtrates from healthy donors can help resolve and protect against infections (13,14) or exacerbate disease phenotypes (15), phenomena potentially mediated by bacteria-phage interactions. Phage particles can also interact directly with the human immune system (16). These results suggest that quantitative characterization of gut phage communities is likely critical for understanding and engineering the gut microbiome.

However, while the individual members of the gut virome are becoming increasingly well characterized, much less is known about their ecological dynamics within a typical human and the effects they exert on the surrounding microbial community. In marine ecosystems, phage particles outnumber bacteria ∼10:1 (17,18) and are estimated to kill ∼20% of the bacterial population each day (18,19). Such high rates of lysis generate strong selection pressures for both bacteria and phage, leading to antagonistic co-evolution (20) and “kill-the-winner” dynamics of strain turnover (21–23). By contrast, estimates of the virus-to-microbe ratio (VMR) in the human gut vary widely across studies, from greater than 1:1 (24,25) to less than 1:10 (26,27). Furthermore, while some studies have suggested that the gut microbiome is dominated by temperate phage (8,28), little is known about rates of induction and lysis, and other studies have suggested that evasion of phage-mediated lysis is a major driver of bacterial evolution within human hosts (11,29). Inferring these ecological parameters is particularly challenging in the complex setting of the human gut, as it requires linking existing measurement approaches with quantitative models of phage population dynamics.

Here, we address this gap by combining mathematical modeling and publicly available data to obtain quantitative baseline estimates of gut viral populations sizes and induction rates in human hosts. Using a meta-analysis of gut viral population size measurements, we show that existing data can be unified into a coherent quantitative picture in which the gut microbiome has more phage genomes than bacterial genomes, but many fewer phage particles than bacterial cells. This suggests that the gut is dominated by temperate phage (here, “temperate” refers to all phage that reproduce with their host genome, including classical lysogens, phage-plasmids, etc.). Building on this quantification, we develop a modeling framework that enables inference of mean gut phage induction rates from microscopy and metagenomic measurements. Our findings suggest that, in typical adults, gut phage are rarely induced and place a low mean fitness burden on their bacterial hosts. Finally, we show that similar ecological dynamics arise in gnotobiotic mice colonized with a synthetic community of >100 human gut bacterial isolates. As expected for a microbiome dominated by temperate phage, we find that the virome composition of these mice can be quantitatively predicted from the bacterial composition alone, while still exhibiting viral diversity comparable to a typical human stool microbiome. These results suggest that existing methods for predicting gut phage lifestyles drastically overestimate the fraction of lytic phage, indicating that many gut phage contain yet uncharacterized host-association genes

## Results

### The typical human gut microbiome contains fewer phage particles than bacterial cells

To determine the range of phage population sizes and virus-to-microbe ratios (VMRs) in the gut, we compiled measurements across multiple methodologies and studies (**Table 1**). Although the VMRs, initially appeared to vary across studies, we found that they could be unified into a coherent quantitative picture by employing a consistent calculation approach that accounts for key differences among existing phage quantification techniques (**Fig. 1A, Methods**).

**Figure 1:**
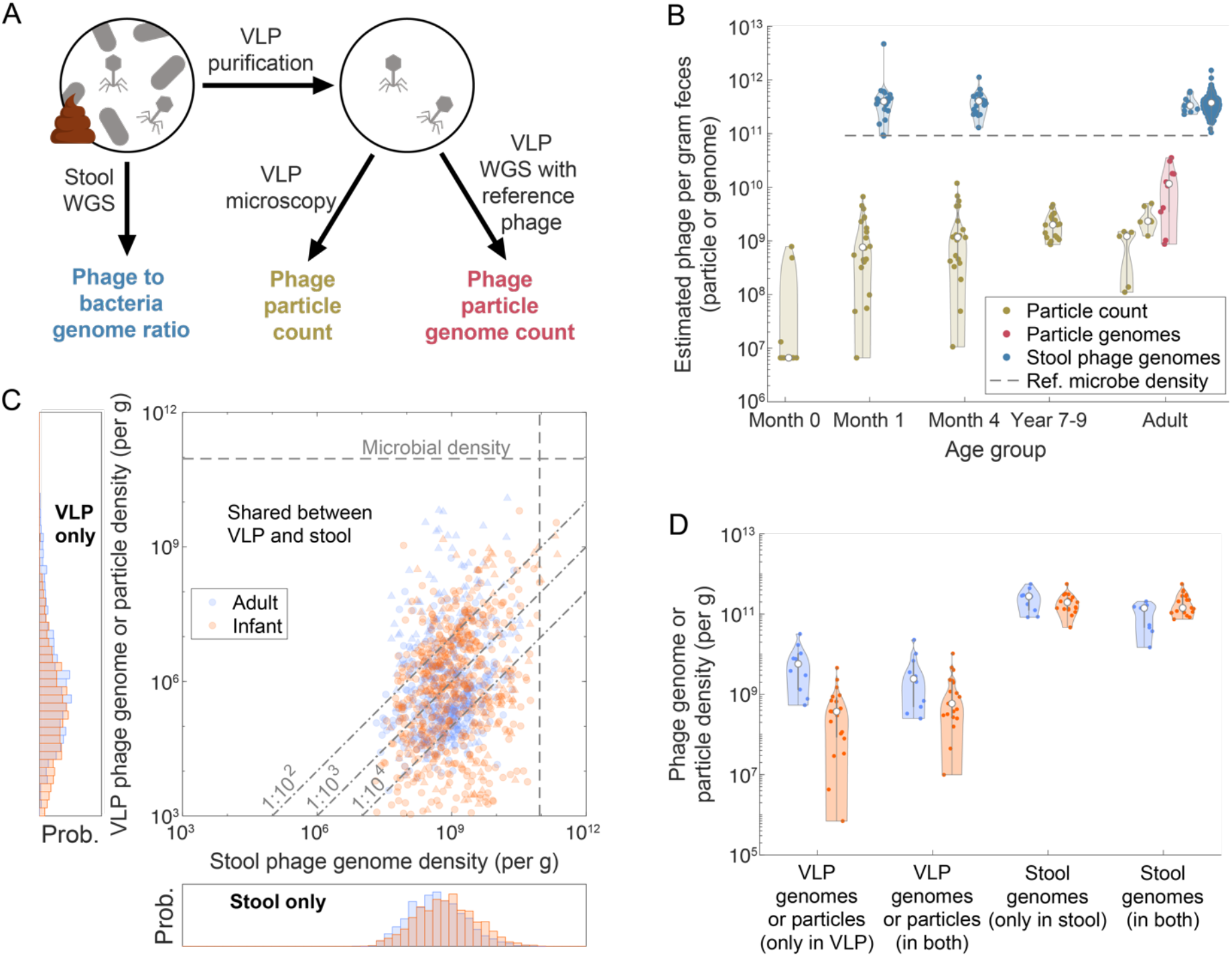
Comparisons of absolute population densities suggest that the human gut virome is numerically dominated by prophage. (A) Schematic overview of common gut viral population quantification methods. Metagenomic classification of bulk fecal samples yields phage-to-bacteria genome ratios, which can be combined with absolute bacterial densities to estimate the absolute density of phage genomes. Alternatively, virus-like particles (VLPs) can be extracted from the stool and quantified by epifluorescence microscopy or spike-in sequencing. (B) Gut virome population densities are approximately maintained across human life stages. Each violin plot represents quantification of one population using one measurement method in one study (**Table 1**), with individual dots representing subjects. The gray line denotes the average stool bacterial density reported in (33) (0.92×10^11^ bacteria/g stool). (C) Species-level absolute abundance analysis of the overlap of phage communities found using VLP- and stool-based quantification approaches. Data represent one healthy adult population (26) (*n* = 10, VLP WGS absolute quantification) and one population of 4-month-old infants (24) (*n* = 19, VLP microscopy absolute quantification). Each point is the absolute abundance of one phage species in a matched pair of VLP and stool samples from one subject (**Methods**). Triangle markers denote species classified as virulent and circle markers denote species classified as temperate (**Methods**). Histograms show the distribution of absolute abundances of phage species found exclusively in either the VLP or stool samples. (D) Relative distribution of phage genomes or particles between VLP and stool WGS. Each violin plot represents the total absolute abundances of phage genomes or particles found within only VLP samples, only stool samples, or shared between VLP and stool. Individual points correspond to a single subject. Underlying data are the same as in (C).

Many approaches for estimating phage abundance involve the isolation of virus-like particles (VLPs) as representatives of the free phage particle population within stool samples. In the most common method, isolated VLPs are enumerated via epifluorescence microscopy using a DNA-binding dye (24,30). These microscopy-based methods estimate the concentration of free phage particles in the stool, although their accuracy is constrained by VLP isolation efficiency (24,31) and the presence of non-phage particles (32). Aggregating VLP enumeration data from multiple studies and age groups (**Fig. 1B**), we found that, apart from newborns in which VLP densities are often below the limit of detection (24), stool VLP density stabilizes after >1 month of age to a population average of ∼2×10^9^ VLPs/g stool, which is maintained throughout adulthood. Combining these data with existing estimates of the density of microbial cells in stool in humans older than >1 month (∼10^11^ cells/g stool; (33)) yields a VLP-to-microbe ratio ∼10^−2^. This estimate is three orders of magnitude lower than the VMRs commonly reported for surface seawater systems (21), hinting qualitatively different viral ecological dynamics that we will explore in more detail below. We also find that the inter-individual variation in VLP counts is similar in magnitude to that of bacterial counts, with post-infancy VLP measurements exhibiting a population coefficient of variation (CV) of 0.61 versus 0.46 for bacterial counts (33). This suggests that the total gut phage population does not undergo dramatic abundance fluctuations across hosts.

An alternative form of phage particle measurement utilizes a spike-in approach, involving shotgun sequencing of amplified DNA from the VLP pool after adding a known amount of a non-gut reference phage (26). The fraction of sequencing reads mapping to reference versus non-reference phage can then be used to obtain an independent estimate of absolute phage particle density (**Fig. 1A**). A recent application of this approach to longitudinal samples from ∼10 healthy adults yielded a ∼5-fold higher concentration than microscopy-based studies (mean of ∼1 × 10^10^ VLPs/g, inter-individual CV of 0.9, **Fig. 1B**). These data also provided an estimate of the temporal variation, with monthly VLP estimates within individuals having a mean CV of 0.78, suggesting that the total phage load in individual hosts does not undergo dramatic fluctuations. The differences between this study and microscopy-based quantifications may be due to underestimation of viral counts by imaging-based approaches relative to sequencing/qPCR-based approaches (34). However, we found that the two measurements are largely consistent after exclusion of reads mapping to the *Microviridae* family of phage (**Fig. 1B**), which are thought to be disproportionately enriched by the multiple displacement amplification (MDA) protocol commonly employed in VLP sequencing (35). Regardless, even the larger VMR estimates resulting from sequencing-based approaches including *Microviridae* reads (∼10^−1^) are still far lower than the ∼10:1 ratios reported for surface seawater (21).

### The typical human gut microbiome contains more phage genomes than phage particles

A third class of quantification methods estimates VMRs directly from metagenomic sequencing of stool samples (25). This approach has been enabled by the recent assembly of large databases of viral and prokaryotic genomes from the human gut (4,5,36), from which >98% of reads from a typical stool sample can be classified using taxonomic profilers like Phanta (25). By normalizing the ratio of phage to bacterial reads with corresponding phage and bacterial genome lengths, one can obtain an independent estimate of the VMR. Applying this approach to a collection of 255 previously sequenced adult gut metagenomes yields an average VMR of ∼4:1 (inter-individual CV = 0.38), corresponding to an absolute density of ∼4 × 10^11^ phage genomes/g after multiplying by the typical bacterial density in Ref. (33). These values are two orders of magnitude higher than the VLP-based estimates above.

The discrepancy between these estimates can be reconciled by the observation that bulk stool metagenomics measures the total number of viral genomes in a stool sample, including those encapsulated in bacterial cells (e.g., as prophage), while VLP-based methods only measure free viral particles. Hence, it is useful to distinguish between two distinct abundance measures: the genomic VMR (gVMR), estimated from bulk stool sequencing, and the particle VMR (pVMR), estimated from VLP-based approaches. The two measures are roughly equivalent in environments like surface seawater where the pVMR is much larger than one (and therefore particles dominate the gVMR). However, they can dramatically diverge in ecosystems like the gut where the pVMR is much less than one. In this case, the ∼100-fold difference between the number of phage particles and phage genomes in the gut suggests that the vast majority of gut phage are temperate or otherwise attached to their bacterial hosts. These temperate phage may not be traditional prophage that are integrated into their host’s genome; many gut phage do not contain recognizable lysogeny-associated genes and thus may utilize other host-associated lifestyles, such as those of phage-plasmids (37).

Consistent with this temperate-dominated picture, we found that the ratio of phage genomes to phage particles is also large for many individual viral species. While MDA amplification biases make precise quantification difficult (35), comparisons between matched VLP and bulk sequencing in infants and adults revealed three broad classes of behavior. Some viral species are observed in the bulk metagenome but not in the associated VLP pool (**Fig. 1C, bottom**). These species account for about half of the total phage abundance in bulk stool samples (**Fig. 1D**) and could represent cryptic (38) or inactive (28) prophage, as well as phage that are poorly amplified by MDA (35). A second set of viral species are observed in VLP sequencing but not in the associated bulk metagenome (**Fig. 1C, left**). These species account for about half of the total phage abundance in the VLP pool, and could reflect both MDA amplification biases (35) as well as phage that are poorly captured by bulk metagenomics. Finally, a third class of viral species is present in both the VLP and bulk metagenomes. Their abundances are broadly consistent with the aggregate particle to genome ratio above, with ∼80% of phage-sample pairs having a ratio below 1:100 (**Fig. 1C, Fig. S1**), and contain a mixture of phage species classified as temperate and purely lytic (**Fig. S2**). These species account for the other half of the VLP and bulk phage populations (**Fig. 1D**). These data suggest that even with limitations imposed by MDA biases, a large fraction of gut phage exhibit a generalized form of temperance, with a small population of viral particles maintained by a much larger number of host-associated viral genomes.

### The phage particle to phage genome ratio provides a lower bound on the rate of phage induction

While population sizes and lifestyles are important aspects of gut ecology, they provide only a static picture of the gut virome and its potential interactions with gut bacteria. To interpret these data and estimate the rates of phage induction and lysis in the human gut, we utilized mechanistic models of phage population dynamics over time (39–41).

We begin by considering a simplified model of phage ecology, which approximates each host gut as a well-mixed ecosystem with mass-action kinetics (**Fig. 2A, Methods**). For a single pair of bacteria and phage, this model can be described by a system of three differential equations for the concentrations of uninfected susceptible bacteria (*S*), infected bacteria or prophage (*P*), and free phage particles (*V*):

**Figure 2:**
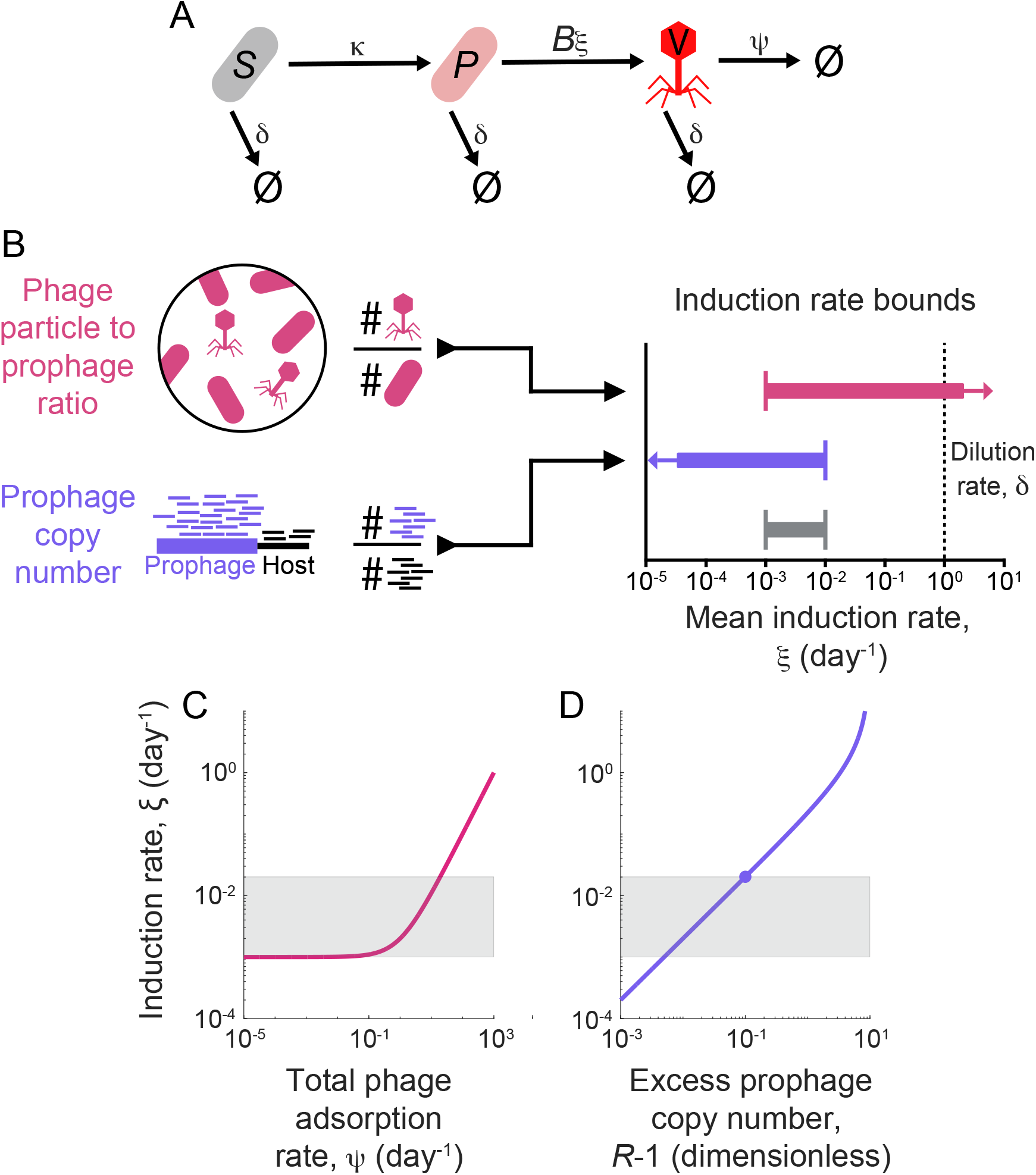
Mathematical modeling of phage population dynamics enables estimation of the average phage induction rate in the human gut. (A) Schematic of the minimal model of temperate phage dynamics represented by **Eq. 1-3**. Infection of susceptible bacteria produces prophage, which induce at rate *ξ*, lysing their host and producing a burst of *B* free phage particles. (B) Schematic representation of induction rate estimates. We combine measurements of phage particle to genome ratio and relative prophage copy number with the model in (A) to estimate upper and lower bounds on the phage induction rate. Note that given the uncertainty in parameter values, these estimates are only reported as approximate orders-of-magnitude, with the combined bound illustrated in grey. (C) Estimated induction rate as a function of total phage adsorption rate ψ (i.e., all non-dilution phage particle removal mechanisms). The solid line corresponds to **Eq. 5**, using the phage genome to particle ratio inferred from **Fig. 1**. (D) Estimated induction rate as a function of the relative coverage of prophage, *R*. The solid line corresponds to **Eq. 8** with *γ* ≈ *δ*. The solid circle is the mean relative coverage in adults (*R* − 1 ≈ 10^−1^), using measurements from (49).

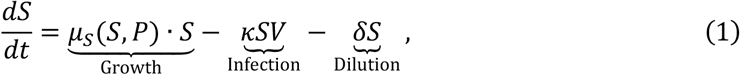

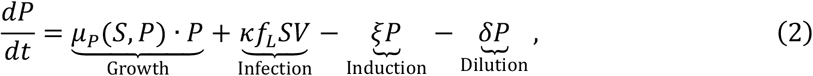

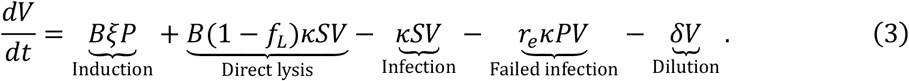

Here, *µ*_*S*_ (*S*, P) and *µ*_*P*_ (*S*, P) are the growth rates of susceptible and infected bacteria, *δ* is the overall dilution rate, and *κ* is the infection rate. We assume that a fraction *f*_*L*_ of infections result in the formation of prophage, while the remaining infections result in direct lysis of the cell with burst size *B*. Phage particles are also produced by induction of prophage at rate *ξ*. We assume that infected cells are immune to further infection by phage particles, with these failed infections resulting in loss of the infecting phage particle (e.g., to superinfection inhibition mechanisms (42)) with rate *r*_*e*_*κ*. We also consider extensions of this model that account for dead cells, dead phage, and actively lysing cells (**Methods**).

Depending on the induction rate and lysogeny fraction, this minimal model can interpolate between a classic lytic lifestyle and a purely temperate phase in which phage primarily reproduce via lysogeny (39,43). In the latter case, the spontaneous induction of prophage can maintain a small population of phage particles (pVMR ≪ 1) while the ratio of phage to microbial genomes (gVMR) remains near one, similar to the distributions seen in **Fig. 1**. This prophage-dominated regime emerges for a broad range of model parameters, particularly when the cost of prophage carriage is low (**Methods**).

We can extend this basic model to larger numbers of phage and bacterial species, except that we must now allow for multiple prophage states in each bacterium (representing simultaneous infection by different combinations of prophage). By summing **Eq. 3** over phage species and integrating over time, one can derive an approximate equation relating the aggregate prophage and phage particle concentrations:

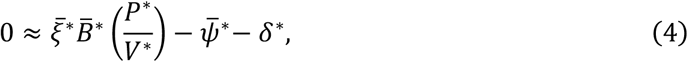

where 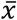 and *x*^*^ denote community- and time-weighted averages of the quantity *x*, respectively (**Methods**), and ψ is the residual phage adsorption rate (e.g., due to failed infections of lysogens). **Eq. 4** assumes that over sufficiently long timescales, the fluxes controlling phage population sizes within an individual (induction, degradation, infection, etc.) are approximately balanced, even though day-to-day fluctuations could still be substantial (we discuss further details of our calculation assumptions in **Methods**). Based on the stability and moderate variance of the distribution of phage population densities (**Fig. 1B**), this assumption appears to hold in healthy humans >1 month of age.

Rearranging **Eq. 4** yields a relation for the average induction rate as a function of the other key model parameters:

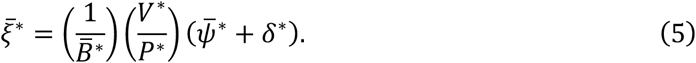

Consistent with intuition, **Eq. 5** predicts that the average induction rate is linearly proportional to the phage particle to genome ratio, *V*^*^/*P*^*^. It also increases with the combined rate of particle removal (dilution rate *δ*^*^ and adsorption rate 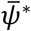, **Fig. 2C**), and decreases with average burst size 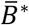, since a smaller number of induction events are required to maintain the same density of phage particles. By combining **Eq. 5** with order-of-magnitude estimates of the other parameters, we can estimate the underlying induction rate. The ratio of phage particle and phage genome densities can be estimated from the population distribution in **Fig. 1** as (*V*^*^/*P*^*^) ≈ 10^−2^. The mean dilution rate *δ*^*^ is determined by the inverse of gut transit time, which can vary across humans but is approximately 1 day^−1^ (44). The burst size *B* can vary substantially across phage, with the model *Escherichia coli* phage *λ* having a burst size *B* ≈ 100 (45), ΦCrAss001 having *B* ≈ L.5 (46), and crAssBcn isolates having *B* ≈ 50 (47). Thus, we estimate the order of magnitude of 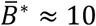. The residual adsorption rate 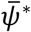 is more difficult to estimate due to our limited understanding of infection rates and host ranges of gut phage *in vivo*. Nonetheless, setting this quantity to zero yields a lower bound on the induction rate,

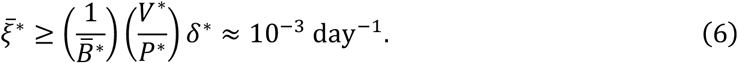

This lower bound increases to 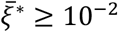 day^-1^ when using the estimate of *V*^*^/*P*^*^ = 10^−1^ from VLP spike-in sequencing but is still two orders of magnitude lower than the dilution rate *δ*^*^. The bound is also relatively tight, with substantial deviations only possible if 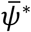 is larger than *δ*^*^ (**Fig. 2C**). These results suggest that the gut phage particle pool, despite having ∼1,000-fold higher density than the highly lytic surface seawater virome (48), can be maintained by a very low rate of induction per infected bacterium.

### The relative coverage of integrated prophage provides an upper bound on the rate of phage induction

Prophage induction can be identified from metagenomic data by comparing the relative coverage of an integrated prophage genome and nearby regions of its bacterial host genome (49). Such methods have thus far been used to make binary determinations of prophage induction for individual phage-bacteria pairs (49), but they also provide information about the underlying induction rate. To extract this information, we use a generalized version of our model in **Eq. 1** to explicitly model activated prophage, representing the state between the start of induction and lysis (**Methods**). These activated lysogens contain *B*_*a*_ ≈ *B* additional copies of the phage genome that correspond to nascent phage particles. The relative coverage *R* of the prophage and host genomes in metagenomic sequencing data is given by

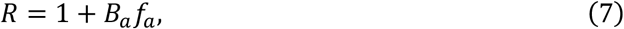

where *f*_*a*_ is the fraction of currently activated cells. If activated cells are produced from lysogens at rate *ξ* and have a mean lysis time of 1/*γ*, then the ratio of activated to non-activated cells will approach a steady-state value of ∼*ξ*/(*γ* + *δ*) (**Methods**). This result can be combined with **Eq. 7** to relate the mean induction rate to the mean relative coverage:

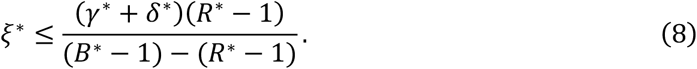

This estimate is robust to the confounding impact of dead cells and viruses contributing to *R* (**Methods**). The relationship between the induction rate and the relative coverage critically depends on the characteristic lysis time of activated infected cell, 1/*γ*. Prior studies suggest that lysis time scales with the bacterial host division time (50–52). This scaling is consistent with estimates of phage burst energetics: a phage burst consumes a large fraction of the host bacterial energy budget (53), implying that production of a phage burst is limited by similar factors as host replication. The mean growth rate roughly matches the dilution rate *δ*^*^ in the parameter regime implied by **Fig. 2C**. Thus, for the following calculations we assume that *γ*^*^ is of the same order of magnitude as *δ*^*^.

**Eq. 8** applies to the subset of phage that are detected within a contig of an assembled bacterial genome. While it in principle enables measurements of arbitrarily low induction rates (**Fig. 2D**), but in practice it is difficult to distinguish small values of *R* from 1 due to noise and biases in sequencing. Indeed, in a previously published analysis of positive and negative controls, induction of individual prophage could only be reliably determined for *R* > L, and the median number of such events across fecal metagenomes was zero (49). To establish a tighter upper bound of the induction rate, we take the “clipped” average of *R* (i.e., setting values of *R* < 1 to 1) across all adult samples analyzed in (49), yielding *R*^*^ − 1 ≈ 10^−1^. Substituting this value into **Eq. 8** with *B*^*^ = 10, and *γ*^*^ ≈ *δ*^*^ = 1 day^−1^ yields an upper bound on the induction rate of

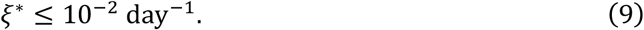

The clipped mean is larger in infants (*R*^*^ − 1 ≈ 0.4), even after excluding infants exposed to antibiotics (**Table 2**), suggesting that the induction rate may be higher in infants. Combined with our other estimates, we can thus bound the average induction rate within the range 10^−3^ − 10^−2^ day^−1^ for adults (**Fig. 2B**), with a somewhat higher upper bound for infants. Importantly, both estimates are substantially lower than the rate of microbial growth and dilution from the gut, suggesting that gut phage impose a low mean fitness burden on their bacterial hosts.

### Similar virome properties arise in gnotobiotic mice colonized with a diverse synthetic community of human gut bacterial isolates

We next examined the implications of our results for a synthetic gut community designed to mimic the complexity of a native human microbiome (54). This synthetic community is composed of 119 bacterial isolates from 48 prevalent genera and stably colonizes gnotobiotic mice for ≥2 months. We reasoned that the virome of hCom2 would be exclusively composed of temperate phage (at least initially), since it was constructed from axenic bacterial cultures (55). Our finding that the human gut is dominated by rarely inducing temperate phage makes two major predictions about the properties of the hCom2 virome and its relation to the human data above.

First, if the induction rates in hCom2 are as low as our model predicts (**Fig. 2**), we expect its viral composition in bulk metagenomic sequencing to be entirely predictable from the abundances of its bacterial members (since *R* ≈ 1). It is usually difficult to test such a prediction in natural communities like the human gut, in which only a subset of phage can be directly linked to their bacterial hosts (49). Synthetic communities like hCom2 provide a unique opportunity to test this prediction, since their initial phage-bacteria associations can be inferred from the sequenced bacterial isolate genomes. To carry out this test, we generated *in silico* hCom2 metagenomes based on data from a recent experiment (54) using the sequenced genome of each bacterial strain in proportion to their measured abundance in each sample (**Methods**). By construction, these *in silico* datasets only contain phage sequences that were present within the original bacterial genomes. We then compared the taxonomic composition of these *in silico* datasets with their corresponding mouse metagenomes using the same pipeline as above (**Fig. 3A,B, Methods**).

**Figure 3:**
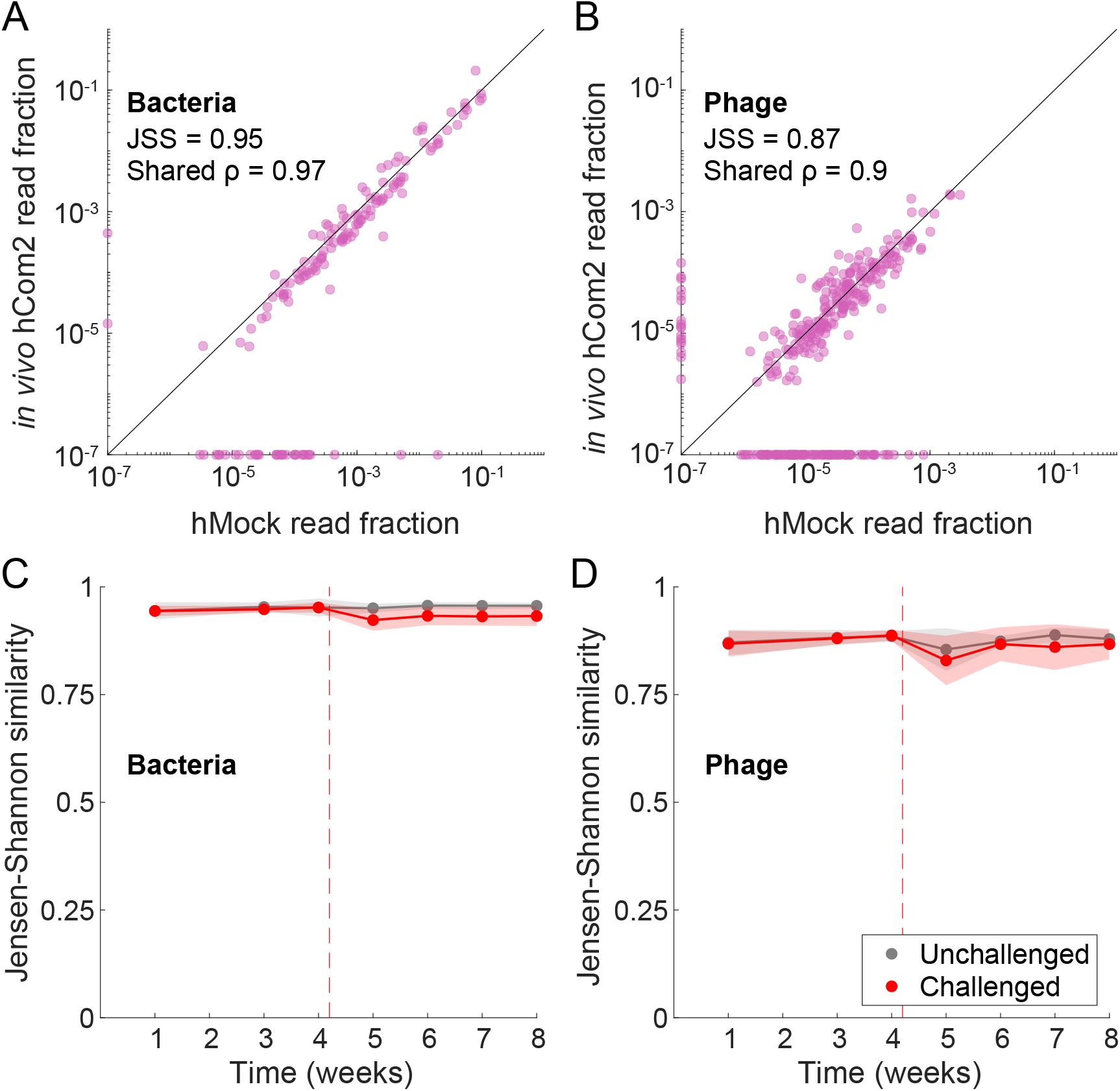
Phage abundance dynamics in a diverse synthetic gut community can be predicted from bacterial abundances alone. (A,B) Relative read abundances of bacterial (A) and phage species (B) in fecal samples from hCom2-colonized gnotobiotic mice, compared to *in silico* metagenomes (“hMock”) generated from their corresponding bacterial genomes weighted according to the fecal bacterial microbiota composition (**Methods**). The example shown is for a single representative sample (mouse 3, week 1). JSS is the Jensen-Shannon similarity and shared *ρ* denotes the Spearman correlation computed from species observed in both samples. (C,D) Bacterial and phage JSS between *in vivo* and *in silico* metagenomes over time and in response to human stool challenge perturbation. Lines show mean JSS in either unchallenged mice (*n* = 5) or mice challenged with a human stool perturbation after week 4 (*n* = 15) over time. Shaded areas represent 1 standard deviation computed across mice at each time point, and the dashed line denotes the time of fecal challenge.

Consistent with previous observations in a smaller 15-member community (8), we found that the abundances of individual phage species were highly correlated across the *in silico* and *in vivo* datasets, with the representative sample in **Fig. 3A,B** having a Spearman correlation of *ρ* = 0.9 for mutually detected phage, compared to *ρ* = 0.97 for bacteria (as expected by construction). Similar results were obtained for other compositional similarity metrics, like the Jaccard index or the total abundance of shared species (**Fig. S3**). The similarity between the *in vivo* and *in silico* metagenomes was maintained over time, and even after challenge with an undefined fecal sample (**Fig. 3C,D, Fig. S3**). These strong correlations confirm that the hCom2 virome is dominated by temperate phage, and that the induction rates are consistent with our inferences from the human data above.

A second – and much stronger – prediction of our prophage-dominated human gut model is that the hCom2 stool virome should qualitatively resemble the stool virome of a typical human. We tested this prediction by comparing the taxonomic composition of hCom2-colonized mouse fecal samples (54) with that of a cohort of 245 healthy human stool metagenomes (56). We reasoned that if hCom2, a synthetic community composed of axenic bacterial cultures, was missing a large portion of the normal gut virome, then feces from hCom2-colonized mice would have substantially lower virome diversity than a typical human stool sample. However, we found that hCom2-colonized mouse feces exhibited similar phage Shannon diversity as human stool samples, with the hCom2 samples falling between the 13^th^ and 53^rd^ percentiles of the observed human distribution (**Fig. 4A**). We obtained a similar correspondence between hCom2 and human stool using a metric of species richness (**Fig. 4B, Fig. S4**), as well as the overall ratio of phage-to-bacterial genomes (**Fig. 4C, Fig. S4**). This similarity between hCom2 and human stool viromes also holds at finer taxonomic levels, with 16 of the 20 most prevalent phage genera within the human cohort found at >0.1% phage community abundance in hCom2 samples (**Methods**). Thus, consistent with our estimates above, we find that human-like levels of viral diversity can be achieved by a synthetic community of exclusively prophage.

**Figure 4:**
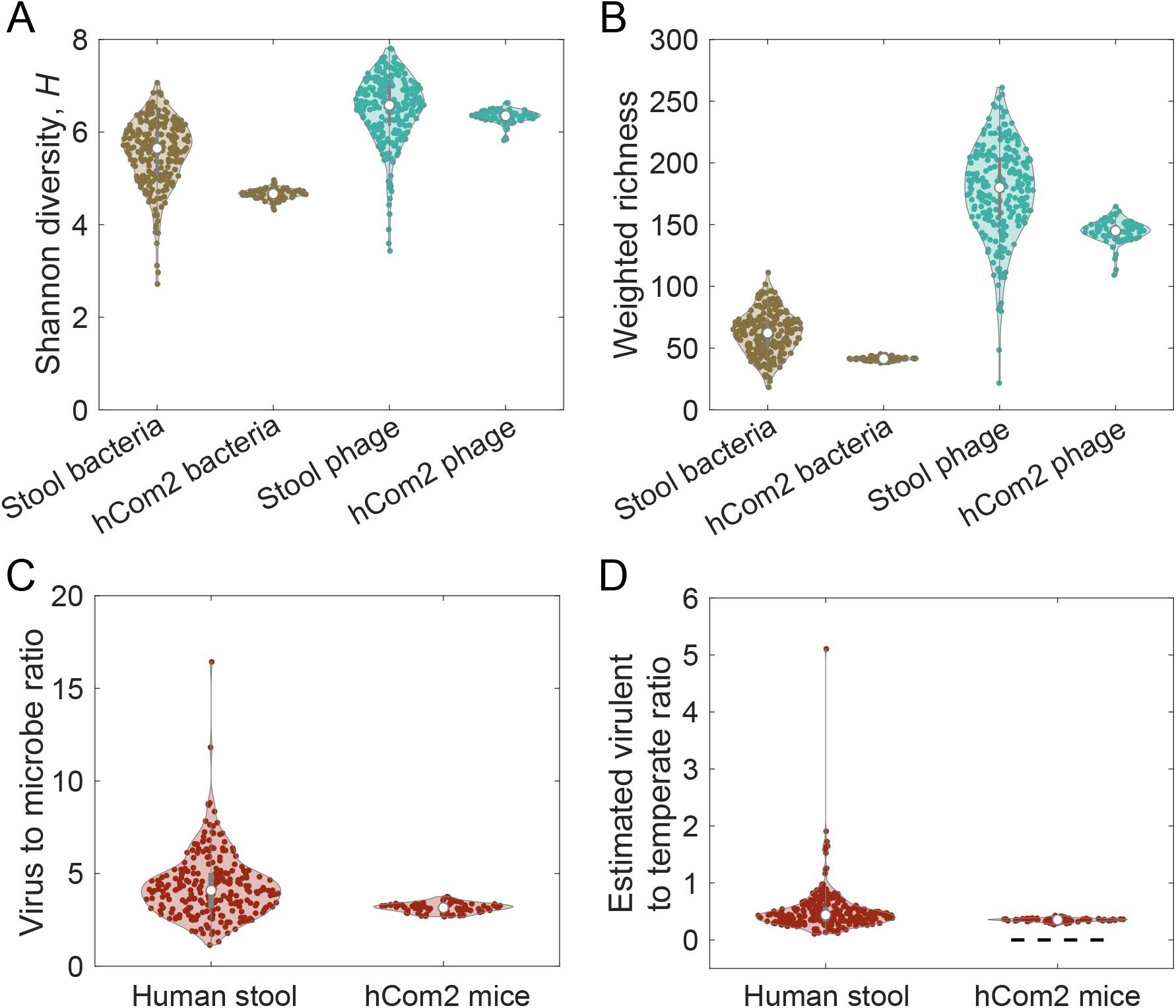
Large-scale features of human stool viromes are recapitulated in a community constructed only of bacterial isolates. (A-D) Comparison of Shannon diversity (A), weighted species richness (B), virus to microbe ratio (C), and virulent to temperate ratio (D) in fecal samples from hCom2-colonized gnotobiotic mice (54) compared to human stool samples. Violin plots labelled “stool” represent distributions of microbiome properties across *n* = 245 healthy adults studied in (56). Violin plots labelled “hCom2” represent samples from 20 gnotobiotic mice colonized with the synthetic community hCom2 (*n* = 77 total samples, all from unchallenged mice or pre-challenge). Virulent to temperate ratios (VTRs) were estimated using the UHGV database phage species virulence predictions (**Methods**). Dashed black line denotes the null expectation of VTR = 0 for a community constructed from axenic bacterial cultures.

The striking similarities between the viromes of hCom2-colonized mouse feces and human stool can shed light on other coarse-grained features of the human gut virome. For example, computational tools have been developed to predict the lifestyles of phage species from their genomes (57,58), enabling estimation of the ratio of virulent phage (those that cannot stably replicate within their hosts) to temperate phage (25,58). We used predictions from widely used tools to estimate the virulent to temperate ratio (VTR) in hCom2-colonized mouse fecal samples (**Methods**). Since hCom2 was constructed entirely from axenic bacterial cultures, it might be expected to provide a negative control with a VTR of ∼0. However, hCom2-colonized mouse feces metagenomes yielded a VTR of ∼0.5, similar to the typical values observed in human stool samples (**Fig. 4D, Fig. S4, Fig. S5**). This result suggests that existing methods of phage lifestyle prediction methods underestimate the number of phage that are capable of stable replication within their bacterial hosts, consistent with previous observations from human stool samples (**Fig. S2**) (26).

## Discussion

Our results complement existing surveys of gut phage diversity (4,24,26,59) by providing a quantitative assessment of phage population dynamics in typical human hosts. Our updated estimates of the virus-to-microbe ratio show that the small number of gut phage particles (pVMR ∼ 10^−2^–10^−1^) is accompanied by a much larger number of phage genomes (gVMR ∼ 4), implying that the vast majority of gut phage genomes are replicating within their bacterial hosts. These results support the emerging view that temperate phage lifestyles play a dominant role in the human gut (8,26,37,60,61), even if they do not contain recognizable integrase genes (e.g. owing to utilization of novel integrases or having non-integrative lifestyles) (26,37) (**Fig. 4D**). Our quantitative framework extends this picture by providing new insights into the corresponding phage induction rates. By integrating imaging and sequencing measurements with a generalized model of temperate phage dynamics, we estimated that the average induction rate in adults lies in the relatively low range of 10^−3^–10^−2^ per bacterium per day, imposing only a modest fitness burden on gut bacteria.

These results starkly contrast with well-studied examples like surface seawater, which possesses a larger ratio of phage particles (pVMR ≈ gVMR∼10) and a higher average lysis rate (21). The reasons for this difference remain uncertain, but they may partially stem from the distinct physical structures of the two ecosystems. Previous theoretical and experimental studies have shown that increased spatial structure can select for lower virulence and increased lysogeny (62,63), consistent with the dominance of temperate phage in the more spatially structured gut ecosystem, although more work is needed to quantify the strength of this effect in the gut. Regardless, our results establish baseline expectations for the co-variation between phage and bacterial abundances within the microbiome of a typical human. They imply that tight associations between the bacterial and phage communities may not be driven by active predator-prey interactions, but may instead be a simple consequence of their synchronized replication within the same cells, in line with the “piggyback-the-winner” model (61,64,65). This latter scenario suggests that phage may impact the gut microbiome primarily by acting as genetic cargo, altering the behavior of their bacterial hosts in certain conditions (11,66).

These results have substantial implications for future studies of the gut virome’s role in human health. Many studies have sought to identify biomarkers and characterize possible mechanistic links between gut virome composition and health states such as lifespan (12), cancer treatment response (2), diabetes (67), metabolic syndrome (68), and alcoholic hepatitis (69). Importantly, our results highlight confounding factors that complicate such analyses of virome-health associations, particularly for studies focused on bulk stool sequencing in which a high abundance of prophage will likely result in strong statistical links between phage and bacterial composition if the number of VLPs is low. In studies focused on VLP sequencing, similar correlations could emerge if the VLP pool is largely a product of relatively uniformly induced prophage, a scenario hypothesized by prior work (24) and supported by the substantial overlap between bulk and VLP virome compositions (**Fig. 1C**) (25). These results suggest that methods similar to phylogenetic regression (70) may be useful for dealing with these confounding factors.

The quantities in our modeling framework represent averages over time, space, and hosts that may mask important behaviors that are transient or localized to a particular host microniche. In particular, averages over longer timescales may not capture shorter-term variation in induction rates. While the VLP population appears to be broadly stable over long timescales, the monthly CV of VLP abundances within individuals has a mean of 0.77 (26). One explanation for such variation is phage induction driven by environmental changes within the host, a hypothesis consistent with prior studies showing increased lytic activity in response to perturbations such as bacterial/phage invasion (8,11), inflammation (71), or exposure to certain dietary or pharmaceutical compounds (72,73). In addition to such temporal and host variation, phage population sizes and induction rates may also vary spatially within an individual gut (61,74–76), as environmental conditions and bacterial densities change substantially along the gastrointestinal tract (77). It remains possible that the low pVMRs observed in stool could be produced by a very high induction rate in a smaller population of bacteria in the proximal colon or small intestine. In the future, spatial variation could be investigated using recently developed methods for spatially resolved sampling of the microbiome (78) to measure population sizes and prophage copy numbers across the intestines. Our modeling framework can be readily applied to such data to estimate local virome induction rates.

Variation in lifestyle characteristics across phage is also expected, with some phage effectively existing as mobile genetic elements that rarely lyse their host and others being primarily lytic. While our current estimates average over multiple phage taxa, our modeling framework can also be applied to measurements of individual phage species to estimate species-specific properties. For example, if the particle-to-genome ratios of an individual phage species can be more accurately measured (**Fig. 1C**), a species-specific estimate of the induction rate can be obtained from **Eq. 5**. Applications of this approach are currently limited by the known amplification biases of existing VLP sequencing methods (35), but the adoption of sequencing protocols that do not involve MDA (79) may enable such species-specific resolution in the future.

Beyond our modeling assumptions, there are also limitations in the phage quantification methods used for experimental measurements. The process of VLP isolation may lead to substantial loss of phage particles, particularly given the spatially structured nature of stool and the potential for phage particles to adhere to large particulates. Additionally, imaging-based quantification methods can both underestimate phage densities due to loss of material during preparation (34), and overestimate due to the presence of cell debris or other DNA-containing particles (32). Similarly, RNA phage cannot be visualized using DNA-staining-based microscopy (24,30). Underestimation of phage particle densities would imply a higher true pVMR, which would increase the corresponding induction rate estimate from **Eq. 5**. Note, however, that for our pVMR estimates to be comparable to that of surface seawater would require very large differential loss rates (>99%), which could potentially be measured with appropriate spike-ins. Our analysis framework can easily be applied to updated density estimates as they become available.

Overall, our work motivates future experimental directions for the gut virome field. While informative, our estimates of the mean induction rate still encompass 1-2 orders of magnitude owing to limitations of current data. Given the noise intrinsic to metagenomic sequencing, we expect that deeper bulk sequencing will have limited benefits for estimating of the mean induction rate in the parameter regimes suggested by our analysis. More accurate and direct estimation will likely be dependent on measurement of rare induced cells. Single-cell bacterial sequencing (80) is a promising avenue to achieve the needed detection power. Alternatively, measurement of *in vivo* phage adsorption rates or the degradation rates of lysed cells (**Methods**) would enable improved estimation of induction rates that we derived in **Fig. 2**. Our results also provide guidance for the design of virome perturbation experiments, which should focus on measuring increases in induction and horizontal gene transfer – a major avenue through which prophage influence their hosts. Finally, the similarities between the estimated VTR in hCom2-colonized mouse feces (**Fig. 4D**) and human stool metagenomes highlights the current lack of knowledge regarding the genetic mechanisms enabling bacterial host-association of gut phage. These results imply that many gut phage currently computationally identified as virulent in fact contain unidentified and uncharacterized host-association genes. This pool of genes represents a rich ground for future phage molecular biology work.

## Acknowledgements

The authors thank the Huang and Good labs, Ami Bhatt, Danica Schmidtke, Gabriel Birzu, Colin Hill, and Andrey Shkoporov for helpful discussions. The authors acknowledge support from NIH RM1 Award GM135102 and R01 AI147023 (to K.C.H.), NSF Awards EF-2125383 and IOS-2032985 (to K.C.H.), NIH R35 GM146949 (to B.H.G.), Alfred P. Sloan Foundation grant FG-2021-15708 (to B.H.G.), Human Frontier Science Program grant RGEC33/2023 (to B.H.G.), and a Friedrich Wilhelm Bessel Award from the Humboldt Foundation (to K.C.H.). B.H.G. and K.C.H. are Chan Zuckerberg Biohub Investigators. J.A.L. was supported by a Stanford PRISM Baker Fellowship. This work was also supported in part by the National Science Foundation under Grant PHYS-1066293 and the hospitality of the Aspen Center for Physics. We thank the Stanford Research Computing Center for use of computational resources on the Sherlock cluster.

## Methods

### Meta-analysis of gut phage quantifications

Table 1 summarizes the studies used in our meta-analysis of gut phage abundances. Each row corresponds to a single violin plot in **Fig. 1A**, with the order of the table rows matching the order in which the datasets appear in the figure.

**Table 1:** Studies represented in the quantification meta-analysis in Fig. 1A. “Data origin” column indicates the study that produced the original data, and “Secondary analysis” denotes studies that performed additional bioinformatic analyses represented in **Fig. 1A**. For all stool WGS quantifications, Phanta was used to estimate the genomic virus-to-microbe ratio (gVMR) which was then multiplied by an estimate of gut microbial abundance (33). For studies in which a table of quantifications was not explicitly provided, counts were digitally extracted from figures using WebPlotDigitizer.

| Data origin | Secondary quantification | Method | Population |
| --- | --- | --- | --- |
| Liang *et al.* 2020 (24) | N/A | VLP EFM | Newborns |
| Liang *et al.* 2020 (24) | N/A | VLP EFM | 1-month-old infants |
| Liang *et al.* 2020 (24) | This study | Stool WGS | 1-month-old infants |
| Liang *et al.* 2020 (24) | N/A | VLP EFM | 4-month-old infants |
| Liang *et al.* 2020 (24) | This study | Stool WGS | 4-month-old infants |
| Bikel *et al.* 2021 (81) | N/A | VLP EFM | 7–10-year-old children |
| Kim *et al.* 2011 (27) | N/A | VLP EFM | Healthy adults |
| Hoyles *et al.* 2014 (30) | N/A | VLP EFM | Healthy adults |
| Shkoporov *et al.* 2019 (26) | N/A | VLP WGS | Healthy adults |
| Shkoporov *et al.* 2019 (26) | This study | Stool WGS | Healthy adults |
| Yachida *et al.* 2019 (56) | This study | Stool WGS | Healthy adults |

We now briefly describe the different measurement methodologies applied by the studies analyzed here. The measurement methodologies for gut phage abundance fall into three classes (**Fig. 1A**), which quantify different subsets of the gut phage population. One method (labeled “Stool WGS” in **Table 1**) is based on metagenomic sequencing of whole stool samples, from which the ratio of the abundance of phage DNA to that of bacteria DNA can be computed (25). By normalizing this ratio by typical phage and bacterial genome lengths, the ratio of phage to bacterial genome copies is obtained (25), which combined with quantification of absolute bacterial density generates an estimate of absolute phage genome density. This method captures both prophage (e.g., lysogens, phage-plasmids, etc.) and the fraction of phage particles that lyse during DNA extraction. The other two methods involve isolation of virus-like particles (VLPs) as representatives of the phage particle population present within stool. Isolation typically involves 0.2-or 0.45-µm filtration and DNAse/RNAse treatment, among other steps. In one method, VLPs are stained with a DNA-binding dye and enumerated via epifluorescence microscopy (24) (labeled “VLP EFM” in **Table 1**), while in the other method the VLPs are mixed with a known quantity of a non-gut reference phage and metagenomically sequenced, with the reference phage enabling absolute quantification (26) (labeled “VLP WGS” in **Table 1**). In contrast to the method based on bulk stool metagenomics, these VLP-based methods do not capture prophage by design. Given the drastic methodological differences between bulk and VLP-based approaches, we define two separate VMRs: the genomic VMR (gVMR), based on bulk stool sequencing, and the particle VMR (pVMR), based on VLP-approaches.

In our calculations of pVMR and of phage absolute abundance from gVMR, we require an estimate of the microbial density of the gut microbiome. Note here that we use the term “microbe” to denote all microorganisms (including archaea, bacteria, and unicellular eukaryotes); in practice, the vast majority of gut microbes are bacteria (33) and this is reflected in our taxonomic estimations from Phanta. For all such calculations, we used a standardized value of 0.92×10^11^ microbes/g stool obtained from a comprehensive meta-analysis of stool microbe abundance quantifications from humans >1 month of age (33). Using a single standardized value is justified by the minimal variation of total gut microbial density across human populations >1 month of age (33). Doing so also eliminates the confounding effect of inter-study variability gut microbial density. Indeed, a few gut virome studies reported bacterial density estimates of ∼10^9^ or ∼10^10^ microbes/g stool (24,27), orders of magnitude below well-established values, which led to inflated values of pVMR. We do not know the origin of these discrepancies, but we assume based on the weight of evidence that the microbial density is closer to 0.92×10^11^ microbes/g stool.

For the direct VLP-stool comparisons in **Fig. 1C,D**, we used only subjects for which VLP metagenomes, stool metagenomes, and VLP absolute quantification were available. For the dataset in (26), matching metagenomes were available for the subjects only at the 8-month timepoint, and VLP quantification was performed only during months 9-12. Thus, we used the month 8 metagenomes in combination with the average of month 9-12 measurements of each subject. We found that in both infant and adults these VLP metagenomic samples had a high gVMR, median ∼10^3^ compared a median gVMR of ∼3-4 found in the corresponding stool samples (as measured by Phanta). This large gVMR difference persists even if *Microviridae* species or all species found only in VLP sample are removed, indicating that the VLP-stool overlap in **Fig. 1C,D** is likely not due to bacterial contamination of the VLP pool.

### Analysis of prophage copy number data

For our estimation of induction rate from prophage copy number, we use results from Kieft et al. (49), which developed a computational tool, PropagAtE, for estimating whether an integrated prophage is active. They applied their tool to several metagenomic sequencing studies and we use the values of *R* estimated by their tool (available in Table S3B of their manuscript). We use only prophage-sample combinations detected as present by their tool and perform additional quality filtering requiring minimum median host and prophage coverage >1, and prophage coverage breadth >0.5. We show the resulting summary statistics across cohorts in **Table 2**.

**Table 2:** Summary statistics of prophage copy number *R* across different metagenomic sequencing cohorts, computed based on results from Kieft et al. (49). “*n* samples” denotes the number of metagenomic sequencing samples in the cohort, “*n* pairs” denotes the total number of prophage-bacterial host pairs identified as present and passing the coverage/breadth requirements in those samples. The clipped mean is the mean computed with values of *R* < 1 set to *R* = 1. The CRC dataset is composed of adults with colorectal adenoma and healthy adults, HeQ is composed of adults with Crohn’s disease and healthy adults, IjazUZ is composed of adults with Crohn’s disease, Infant (non-abx) is composed infants that were not exposed to antibiotics, Infant (abx) is composed of infants that were exposed to antibiotics.

| Sample set | $n$<br>samples | $n$<br>pairs | Mean $R$ | Median $R$ | Clipped<br>mean $R$ |
| --- | --- | --- | --- | --- | --- |
| All adults | 123 | 3,459 | 1.04 | 1.01 | 1.09 |
| CRC | 15 | 474 | 1.02 | 0.99 | 1.07 |
| HeQ | 96 | 2842 | 1.05 | 1.02 | 1.09 |
| IjazUZ | 12 | 143 | 1.04 | 1.04 | 1.08 |
| All infants | 79 | 702 | 1.27 | 1.1 | 1.40 |
| Infant (non-abx) | 22 | 254 | 1.33 | 1.22 | 1.43 |
| Infant (abx) | 57 | 448 | 1.24 | 1.02 | 1.39 |

### Processing and analysis of metagenomic datasets

All metagenomic datasets analyzed in this manuscript were first subjected to quality control/filtering and adapter removal using the BBDuk decontamination tool in BBTools (82). Settings used were kmer length “k = 23”, “hdist = 1”, trim direction “qtrim = rl” (trim both ends), minimum entropy “entropy = 0.5”, sliding window for entropy calculation “entropywindow = 50”, kmer length for entropy calculation “entropyk = 5”, minimum quality “trimq = 25”, and minimum read length “minlen = 50”. Samples were then deduplicated using the clumplify tool in BBTools. The maximum number of substitutions between duplicate reads was zero (“subs = 0”). We found that deduplication minimally influenced the estimated community compositions.

For taxonomic quantification of samples, we used Phanta, a kmer-based method that simultaneously profiles phage and bacteria (25). Phanta was run using default settings: confidence threshold “confidence_threshold 0.1”, viral genome coverage requirement “cov_thresh_viral 0.1”, viral unique minimizer threshold “minimizer_thresh_viral 0”, bacterial genome coverage requirement “cov_thresh_bacterial 0.01”, and bacterial unique minimizer threshold “minimizer_thresh_bacterial 0”. The “uhggv2_uhgv_mqplus_v1” database was used, which is based on the prokaryotic UHGG database and viral UHGV database. For taxonomy-based analyses, the provided UHGV taxonomy was used, except for quantification of *Microviridae* abundance, for which we used the provided ICTV taxonomy.

### hCom2 metagenome reconstruction from bacterial genomes

To generate mock versions of hCom2-colonized mouse fecal metagenomes, we generated synthetic short-read sequencing datasets using the set of isolate genomes (54). To determine the relative abundances of genome reads within each sample, we used the bacterial compositions estimated by NinjaMap (54). NinjaMap is designed to quantify the composition of synthetic communities in which sequenced genomes are available for all member strains. For each sample, we specified the relative fraction of reads from each genome based on that strain’s relative abundance and normalized by its genome length. For genomes that are not assembled into a single contig, the read abundance was split among the contigs weighted by the length of each contig. To generate synthetic shotgun samples, we used Grinder (83) with the following settings: quality levels “-qual levels 33 31”, insert distance “-insert_dist 800”, read length “-read_dist 140”, forward-reverse mate orientation “-mate_orientation FR”, characters deleted from reference sequences “-delete_chars ‘-∼*NX’”, and distribution of mutations “-mutation_dist uniform 0”. The total number of reads generated for each sample was equal to the post-QC read number of the corresponding original mouse fecal sample. Fecal samples with <10^5^ reads were excluded from the analysis. The resulting samples were subjected to the standard pre-processing pipeline applied to all other metagenomic sequencing data in the manuscript. For the comparison of hCom2 stool to human stool, we exclude human stool samples with <10^5^ reads.

### Computation of community summary statistics in metagenomic samples

For all analyses except the hCom-hMock comparison in **Fig. 3** and **Fig. S3**, we used relative taxonomic abundances (which are computed in Phanta by normalizing relative read abundances to the phage or bacterial genome size). For the hCom-hMock comparison, relative read abundances were used to better compare reconstruction fidelity between phage and bacterial communities. To obtain the genomic virus to microbe ratio (gVMR) of a sample, we calculated the ratio of the total taxonomic abundance of members within the viral superkingdom to the total taxonomic abundance of members of the archaeal, bacterial, and non-human eukaryotic superkingdoms. In practice, the denominator of the gVMR is vastly dominated by the bacterial taxonomic abundance. To obtain the virulent to temperate ratio (VTR), we calculated the ratio of total taxonomic abundance of phage classified as virulent to the total taxonomic abundance of phage classified as temperate. In the main figures, we used the virulence predictions from the Phanta UHGV database (25), which utilizes a combination of scores from BACPHLIP (57) along with information from the PHROG database (84) and geNomad (85). An alternative VTR estimate was performed with scores from PhaTYP (58) (**Fig. S5**). PhaTYP was run on the UHGV genomes using default settings.

Shannon diversity was computed at the species level as *H* = − ∑_*i*_ *x*_*i*_ log_2_(*x*_*i*_), where *x*_*i*_ is the relative taxonomic abundance of species *i* within the bacterial or phage community. Weighted richness was computed such that the richness contribution of each species is weighted by 1 − exp(−*x*_*i*_/*x*_0_), where *x*_0_ = 10^−3^. For the hCom2 reconstruction analysis, the Jensen-Shannon similarity was computed as 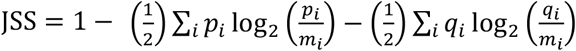, where *p*_*i*_ and *q*_*i*_ are the relative read abundances of the communities being compared, normalized to sum to 1 within a given taxonomic grouping (e.g., phage at the species level), and 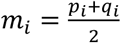.

### Overview of phage mathematical model

We begin with a mathematical model of a single phage and bacterial species in a well-mixed environment, similar to that of (39). We show here how this model can be reduced to the model presented in the main text. This model involves the concentration of a nutrient (*C*), susceptible cells (*S*), cells containing quiescent prophage (*P*), cells in which the prophage has been activated (*P*_*a*_), viral particles (*V*), and dead cells/viruses of various kinds (*D*_*i*_). All populations are diluted at rate *δ*. All populations also experience non-dilution mortality/degradation at rate *ω*_*i*_. All bacterial cells experience the same non-dilution mortality rate *ω*_*B*_. Susceptible cells and cells containing quiescent prophage grow by consuming the resource. Resource consumption occurs with uptake rate *µ*(*C*) for susceptible cells and *µ*(*C*)(1 + *s*) for prophage-containing cells, where *µ*(·) is the growth function and *s* is the fitness benefit/cost of carrying a quiescent prophage. Resources are supplied at a constant rate Γ. Susceptible cells are exposed to viral particle infection by mass-action kinetics at rate *κ*, with a fraction *f*_*L*_ becoming quiescent prophage-containing cells and a fraction 1 − *f*_*L*_ shifting to the activated cell class (*f*_*L*_ models the lysis-lysogeny decision upon initial infection). Prophage-containing cells are induced at rate *ξ*, shifting to the activated class. Cells in the activated class are assumed not to grow and lyse at rate *γ*, producing a burst of *B* viral particles. Viral particles are lost by infecting susceptible cells, failed infection of prophage-containing cells (e.g., to superinfection inhibition mechanisms (42)), and non-dilution mortality. Failed infection occurs at rate *r*_*e*_*κ*, where *r*_*e*_ is the ratio of infection coefficients of prophage-containing and susceptible cells. Dead susceptible, prophage-containing, and activated cells, with concentrations *D*_*S*_, *D*_*P*_, and *D*_*a*_, respectively, and dead viruses with concentration *D*_*V*_, are produced by non-dilution mortality. Cells that die by phage lysis are tracked separately with concentration *D*_*L*_. Non-lysed dead cells are degraded at rate *ω*_*D*_, while lysed cells are degraded at rate *ω*_*DL*_, and dead viruses are degraded at rate *ω*_*VD*_. The dynamics governing this model are thus:

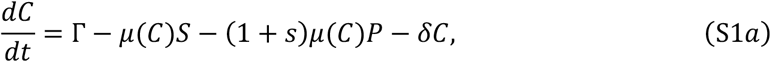

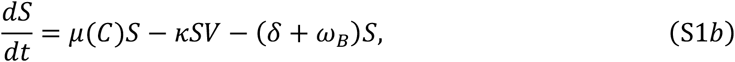

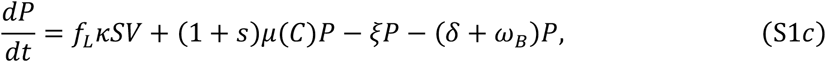

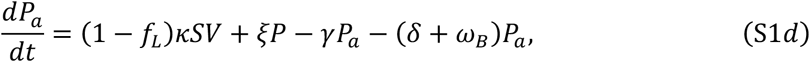

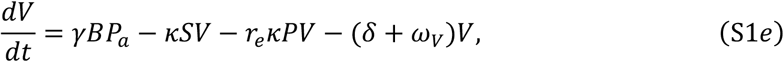

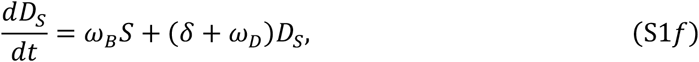

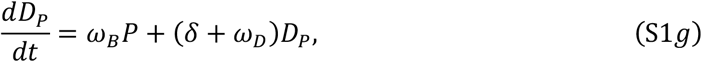

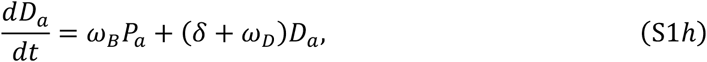

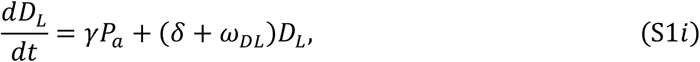

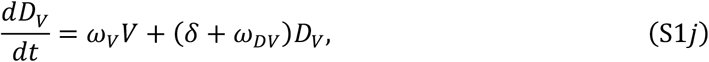

To recover the model discussed in the main text (**Eq. 1-3**), we make a separation of timescales assumption to reduce the number of state variables in the model, assuming that the nutrient, activated cells, and dead cells are in pseudo-steady-state with the remaining state variables (i.e., 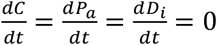). This assumption yields the following expressions:

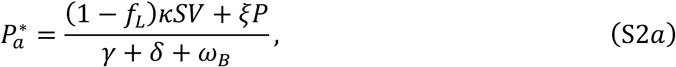

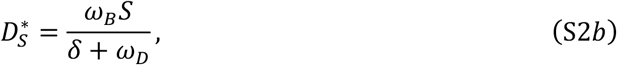

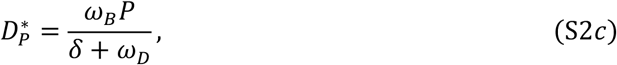

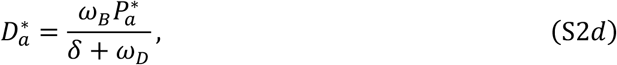

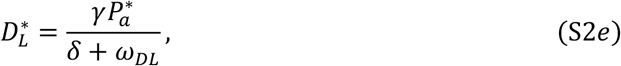

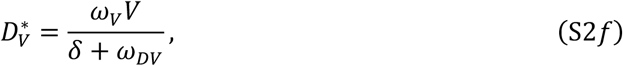

These equations can be used to define a simplified set of equations with only the sensitive, prophage, and phage particle abundances

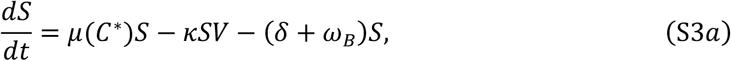

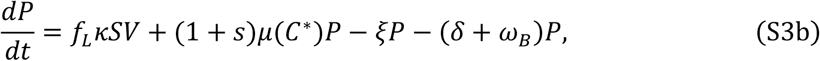

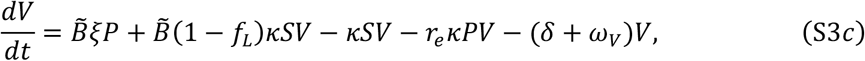

where *C*^*^(*S*, P) is defined implicitly by 0 = Γ − *µ*(*C*^*^)*S* − (1 + *s*)*µ*(*C*^*^)*P* − *δC*^*^ and 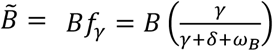. *f*_*γ*_ can be interpreted as the fraction of activated cells that are not diluted or die before lysis occurs and thus 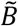 can be interpreted as an effective burst size.

### Conditions for robust phage invasion

In the following two sections, we will assess invasion and stability of phage populations in the model defined by **Eq. S3**. We assume a linear growth function *µ*(*C*) = *αC* for these derivations, leading to 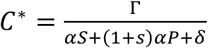. In the absence of bacteria, the resource concentration will saturate at a steady-state value of 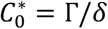. Bacteria will be able to invade this ecosystem when their initial growth rate exceeds the death and dilution rate: 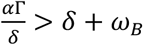. Given the stable bacterial colonization seen in the human gut, we assume this condition to be satisfied. More strongly, given that Γ and *δ* likely vary substantially over time even within a single host (corresponding to variation in food intake and passage time), robust colonization requires 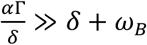.

In the absence of virus, susceptible bacteria will saturate at an equilibrium abundance

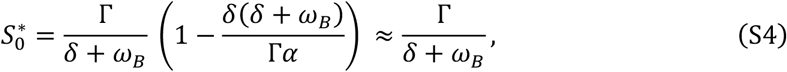

where the approximation follows from the robust bacterial colonization assumption (*α*Γ ≫ *δ*(*δ* + *ω*_*B*_)). In the absence of lysogeny (*f*_*L*_ = 0), viruses will be able to invade this susceptible population if the initial phage replication is greater than death: 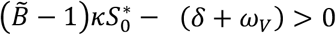. Equivalently, the (lytic) basic reproductive number of the virus must be greater than one:

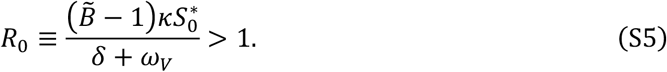

As above, since Γ and *δ* will vary (and thus 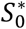 will vary), robust phage invasion will require that *R*_0_ ≫ 1, and thus this is the regime we are primarily interested in.

### Stability of the prophage-dominated steady state

Given the estimated abundance of prophage in the gut (**Fig. 1B**), we are particularly interested in the properties of the prophage-only steady state of the model. We will show that under reasonable assumptions this steady state is likely stable in the gut and thus can be invoked in interpretating our induction rate estimates. The prophage-only steady state has 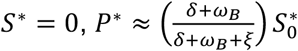, and 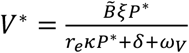, and 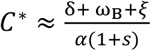. As we are in the robust bacterial colonization regime, we neglect the contribution of dilution to nutrient elimination. This steady state is robust to small perturbations of *P, V*, and *C*. From an invasion analysis, it will be robust to small invasion of susceptible bacteria if the net growth rate of these susceptible bacteria is negative:

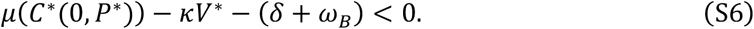

Substituting in the definition of *C*^*^, dividing by *δ* + *ω*_*B*_, and rearranging yields

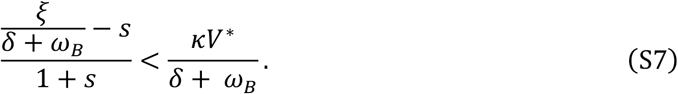

We can express *V*^*^ in terms of *R*_0_ as

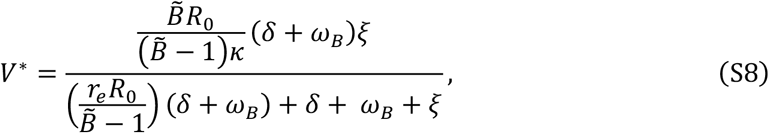

and substituting this equation into the invasion condition yields

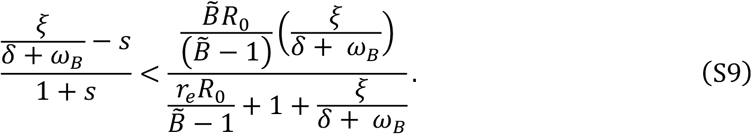

We are particularly interested in the regime where the direct cost (if negative) of the prophage *s* is small relative to one, but still potentially large relative to other small parameters in the system. This limit is consistent with the modest energetic cost of replicating a phage genome (53) and that non-lytic mobile genetic elements have been observed to rapidly undergo compensatory adaptation to reach very low fitness costs (86). Expanding to lowest order in *s* leads to

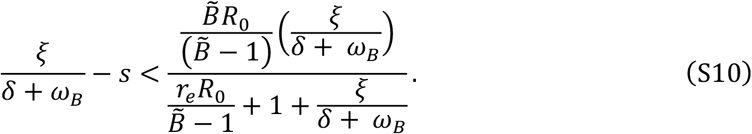

This condition is violated at very high induction rate (*ξ* ≳ *R*_0_(*δ* + *ω*_*B*_)) and at low induction rate when

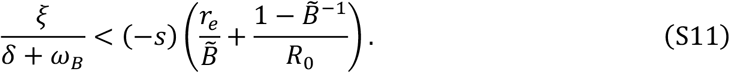

The term on the right-hand-side is much smaller than −*s* in the empirically relevant regime where 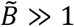 and *R*_0_ ≫ 1. Thus, as long as the dominant cost of gut prophage is induction, i.e., *ξ* > −*s*, as has been experimentally observed for some phage (87), then the gut ecosystem likely exists within a regime where the prophage-only state is stable.

### Overview of induction rate estimation approach

In the following sections, we show detailed derivations of the induction rate estimates presented in the main text, starting from the single phage-bacteria model in **Eq. S3**. In addition to the estimations based on phage particle to prophage ratio and prophage copy number, we also show an estimation based on cell viability. With currently available data, this estimator is poorly constrained and thus not included in the main text, but some results from this derivation are used in the derivation of the estimate based on prophage copy number.

In deriving these estimates, we begin with a general form of the calculation that makes no assumptions about the relative abundance of prophage. This approach leads to estimates of the total lysis rate, which includes both induction of prophage and lysis of sensitive cells via non-lysogenic infection. We then simplify these estimates by assuming that the gut is prophage-dominated, leading to the expressions for the average induction rate in the main text. This simplification only affects the interpretation of the resulting estimate: if the prophage-dominated simplification is incorrect and a substantial amount of phage particle production occurs from sensitive cells, then the estimates are still valid as total lysis rate estimates. In the final model section, we show how our framework can be extended to communities with multiple species of bacteria and phage with explicitly time-varying parameters.

### Total lysis rate and induction rate estimate from phage particle to prophage ratio

Here, we estimate the average lysis rate (including both induction of lysogens and lysis after non-lysogenic infection) from the phage particle to prophage ratio. We begin with the viral dynamics from the timescale-separated prophage model (**Eq. S3**). We define the population-weighted total lysis rate *η* such that *η*(*S* + P) = *ξP* + (1 − *f*_*L*_)*κSV*. We can also rewrite this as *η* = *ξx*_1_ + (1 − *f*_*L*_)*κVx*_*S*_ where *x*_*i*_ are the population relative abundance within the *S* + P pool of cells. By rearranging **Eq. S3c** when 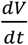 is on average zero (i.e. 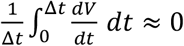), we can obtain an expression for *η*^*^:

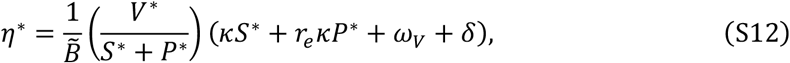

where asterisks denote the time-averaged value 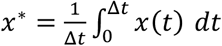. This approximation assumes that the microbiome is in a statistical steady state (no net trend in *V*) and that Δ*t* is long enough that time averages have converged to their ensemble-averaged values. At a minimum, this assumption requires that Δ*t* ≳ 1/*δ*. The assumption of a statistical steady state is supported by the results of our absolute abundance meta-analysis (**Fig. 1**). Given the limited knowledge of *κ, r*_*e*_, and *ω*_*V*_ in the gut, we use **Eq. S12** to construct a lower bound on *η*^*^:

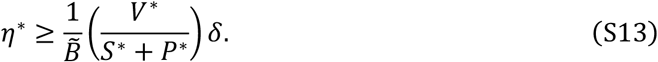

In the prophage-dominated regime (i.e., *S*^*^ = 0) we recover 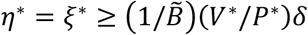, a bound on the induction rate.

In practice, measured pVMR may not be 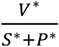 due to the contribution of dead cells and dead viruses. If all populations are represented in the measurement, the pVMR will instead be

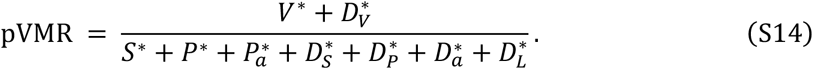

This is related to 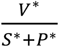 by:

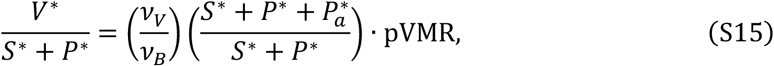

where *ν*_*i*_ is the cell or viral viability fraction (the fraction of cells or viruses that are viable). We now substitute this expression into **Eq. S12** and use the fact that 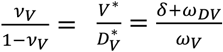 to yield

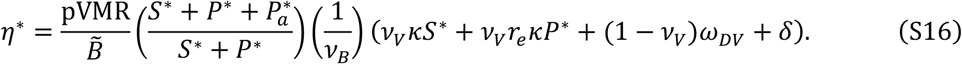

Thus, utilizing a pVMR including the dead populations still functions as a lower bound estimate of *η*^*\**^:

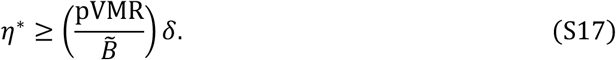

### Cell death and total lysis rate estimates from live cell fraction

Here, we derive estimates of both the non-lysis cell mortality rate *ω*_B_ and the total lysis rate *η* based on the fraction of cells that are living/viable within the microbiome, defined in our model as 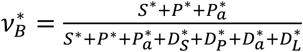. We begin by substituting in the steady-state population abundances to the expression 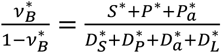

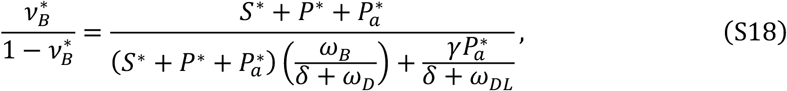

which when solved for *ω*_B_ yields

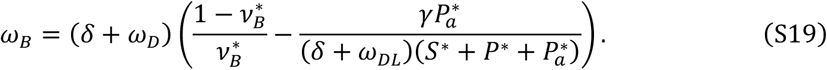

This equation provides an upper bound estimate for *ω*_B_:

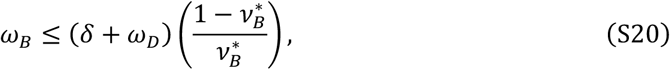

which will be utilized later in deriving the estimate of total lysis rate from prophage copy number. The value of 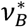 in stool has been estimated at ∼0.5 ™ 0.8 based on cell permeability measurements, leading to 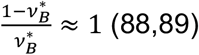. Thus, the bacterial death rate is at most similar in magnitude to the sum of dilution and cell degradation rate.

From **Eq. S18**, we can also derive an estimate for the total lysis rate using the steady-state expression 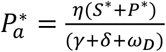. Solving **Eq. S18** for *η* yields:

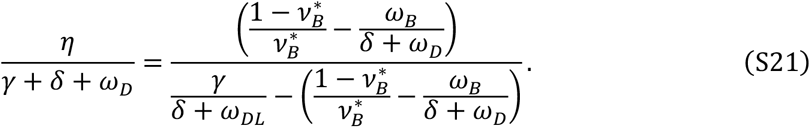

This equation is potentially usable to provide another lysis rate estimate, and in the prophage-dominated regime becomes an induction rate estimate. However, the values of *ω*_D_, *ω*_B_, and *ω*_D*L*_ are currently poorly constrained. For example, one cell viability measurement method is based on comparing the fraction of 16S rDNA found inside and outside of intact cells (88) and it is not known how rapidly extracellular DNA is degraded inside the gut. There are also potential technical issues in the measurement of *ν*_*B*_, as it is unclear to what extent cells lysed by phage are detected by current cell viability measurements. If the lysis process degrades the host genome or leads to total destruction of the cellular structure, the population of cells dying due to lysis would be underestimated by methods relying on extracellular genomic DNA or permeable cell remains.

### Total lysis rate and induction rate estimates from integrated prophage copy number

Here, we estimate the average total lysis rate using lysogeny copy number *R*. We assume that *R* includes the contribution of viral particles, dead viral particles, and all dead cells, and we show that the inclusion of these classes does not substantially alter our induction rate estimation. Each activated cell contributes *B*_a_ prophage copies, each lysogen cell contributes one prophage copy, and lysed cells contribute no prophage copies. All cells contribute a single bacteria genome copy. We first define *R* in terms of our steady-state model populations:

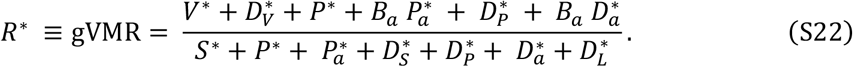

As all cells have the same non-lysis mortality rate, **Eq. S22** can be rearranged to

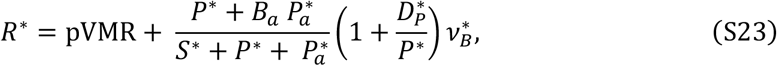

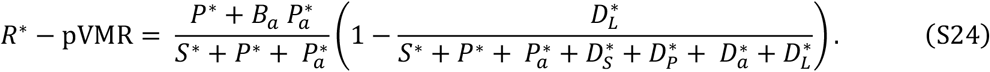

We can then substitute in the steady-state population values to express all dead cell populations in terms of living cells populations:

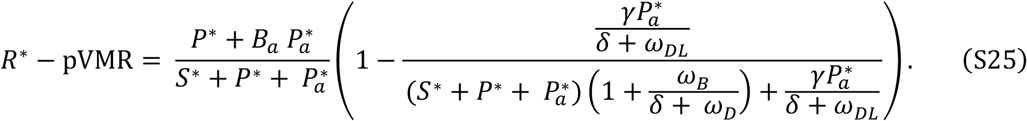

Rearranging and using the fact that 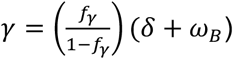 yields

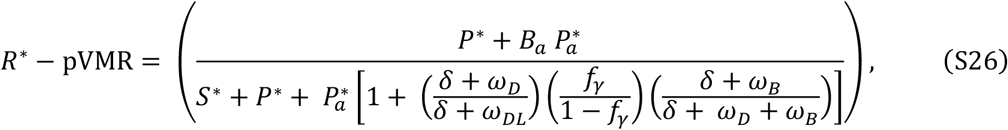

which when solved for 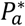 yields

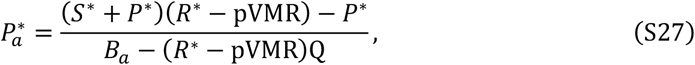

Where 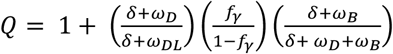 From our steady-state solution for 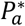 we have that 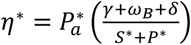, providing an estimate of *η*^*\**^:

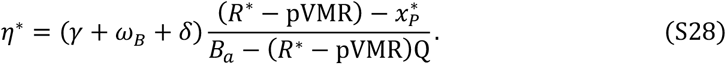

The effect of dead cells and viruses enters the expression via the factor Q, which will inflate the lysis rate. However, this term cannot be greater than *O*(1), and thus if *B*_a_ is large, the impact of dead material is minimal. Empirically, the value of *ω*_B_ is poorly constrained, but we can use results from the cell viability derivation above (**Eq. S20**) to relate this rate to the cell viability fraction *ν*_*B*_ and the degradation rate of dead cells *ω*_D_:

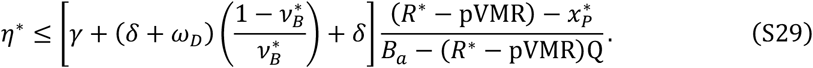

To reach the order of magnitude bound shown in the main text, we assume prophage dominance 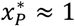, that the number of prophage copies in activated cells is similar to the burst size 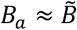, and that dead cells are primarily removed by dilution *ω*_D_ ≪ *δ*. Based on empirical measurements, we also assume that *R*^*\**^ ≫ pVMR,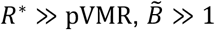, and 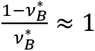, yielding

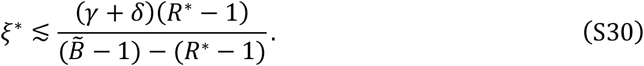

We now briefly discuss potential bioinformatic/sequencing technical artifacts that could influence the measurement of *R*. One potential factor that could systematically skew the above induction rate estimate is sequencing bias between prophage and host (e.g., due to GC content differences between host bacteria and prophage (90)). However, based on the negative control analyses performed by (49), these biases do not appear significant, as non-induced prophage had *R* ≈ 1. If large biases existed, *R* in non-induced phage would differ significantly from 1. Another possible confounding factor in estimating *R* is the presence of the prophage within only a subpopulation of the bacterial host, leading to a lower value of *R*. However, this is unlikely to affect our current analyses, as the *R* values we analyze were computed based on metagenomically assembled contigs containing both prophage and bacterial host sequence. Assembly of such mixed contigs from a mixed lysogen/sensitive population is highly unlikely due to degeneracies in the possible assembly paths. In the case of both possible biases, our framework can readily accommodate improved estimates of *R* as sequencing and bioinformatic approaches improve.

### Extension of the model to multiple phage and bacterial species in time-varying environments

We now generalize our model to complex communities in time-varying environments. For simplicity, we begin with the timescale-separated version of the model and focus on the prophage-dominated case in which most lysis is due to induction, as for the single bacteria-phage regime studied above.

We now track the dynamics of multiple types of phage (indexed by *i*) and multiple types of bacteria (indexed by *j*), such that the total number of phage particles is *V*(*t*) ≡ ∑_*i*_ *V*_*i*_(*t*) and the total number of bacteria is *N*(*t*) ≡ ∑_j_ *N*_j_(*t*). The bacterial “type” *j* encompasses both the taxonomic identity of a bacteria and its infection status (i.e., the *N*_j_ also include bacteria infected by a prophage). To keep track of infection status and the phage-bacteria interaction network, we introduce bookkeeping parameters *I*_*ij*_ and *A*_*ijk*_, respectively. The parameter *I*_*ij*_ is 1 if bacteria *j* is infected with a prophage of phage *i* and zero otherwise. Thus, the total number of prophage in this system is *P*(*t*) ≡ ∑_*i*_ *P*_*i*_(*t*) = ∑_*ij*_ *I*_*ij*_*N*_j_(*t*). Note that generally *P*(*t*) ≠ *N*(*t*), as multiple phage can infect a single bacteria. The second parameter, *A*_*ijk*_, is 1 if an infection of bacteria of type *j* by phage *i* produces an infected bacterium of type *k*, and 0 otherwise. Using this notation, we now define the multispecies generalization of **Eq. S3**:

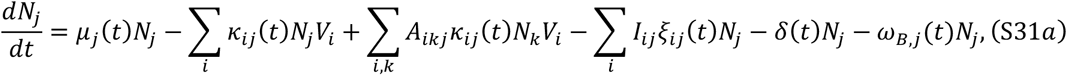

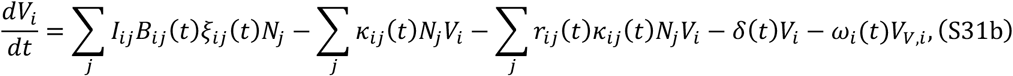

where we have also allowed the rate parameters to explicitly depend on time. We have also approximated *f*_L_ = 1 for simplicity.

To relate these dynamics to the total pVMR, we now sum **Eq. S13b** over the viral index *i* and substitute 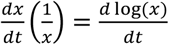, yielding

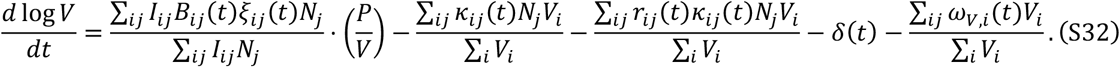

**Eq. S14** can be rewritten in a more compact form as

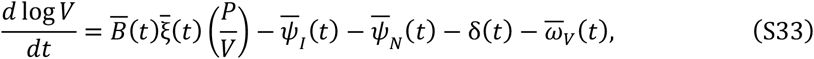

where we have defined the microbiome averages

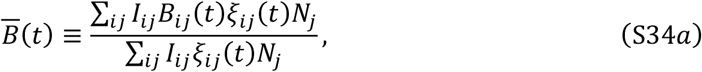

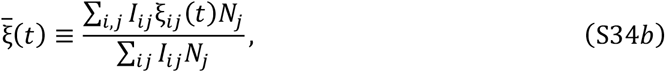

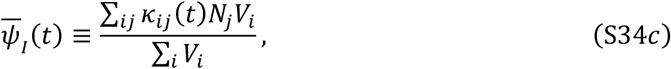

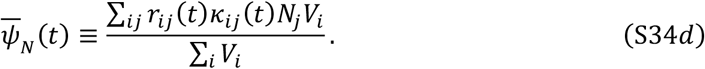

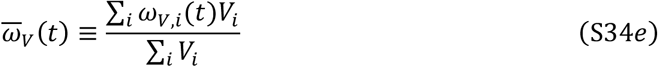

Integrating **Eq. S15** over long times yields

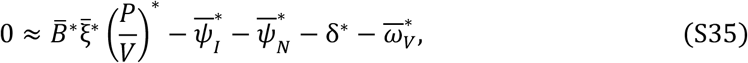

where the asterisks again denote the time-averaged value 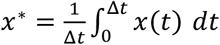. Since the microbiome is in a statistical steady state over long times (**Fig. 1B**), we can estimate the averages over time by taking an average over independent hosts. This procedure yields a connection between the rate parameters and the VLP-to-prophage ratio from **Fig. 1** and thus a lower bound similar to the one estimated from the single phage-bacteria model:

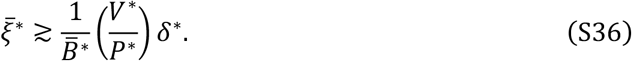

This bound assumes that the burst size, induction rate, and particle-to-prophage ratio are largely uncorrelated in time. If this assumption is violated, the estimate represents a particular weighted average of the induction rate bound:

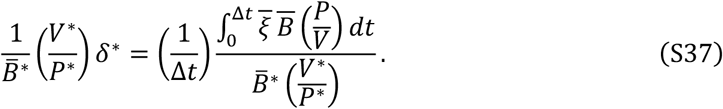

To generalize this bound to the case of pVMR including dead material, we begin with the multispecies version of the dead virus dynamics:

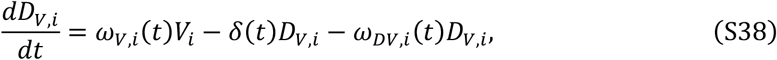

which then yields an expression for the dynamics of the total dead virus population:

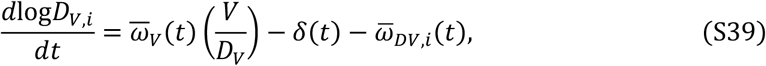

where 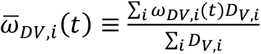. At statistical steady state, this leads to 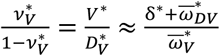 which when combined with **Eq. S35** shows the lower bound is preserved when the pVMR accounts for dead populations, as in the earlier single species derivation.

Note that in this section we have only analyzed a simple case of this community model, and further analysis, such as exploring the role of temporal correlations and the relative contribution of induction and direct lysis, is a promising direction for future theoretical phage ecology work.

Here, we have shown the multispecies generalization of the induction rate estimate from VLP-to-prophage ratio. The rate estimates computed from cell viability will similarly extend to the multispecies context, as we model all sources of death in aggregate, independent of which phage causes lysis. The induction rate estimate from the prophage copy number is performed on a prophage-by-prophage basis, hence it is not affected in a multispecies context. However, this context will lead to a difference in the kind of average used in the estimate: unlike the average computed from the VLP-to-prophage ratio, the average from copy number average is not abundance weighted and includes only lysogens captured with their host contig.

## Data Availability

All code used in this manuscript is available at https://github.com/jamie-alc-lopez/gut_phage_quantification. All data analyzed in this manuscript is publicly available. Processed final versions of the datasets (e.g. estimated taxonomic compositions) are available in the GitHub repository.

## Supplementary Figures

**Figure S1:**
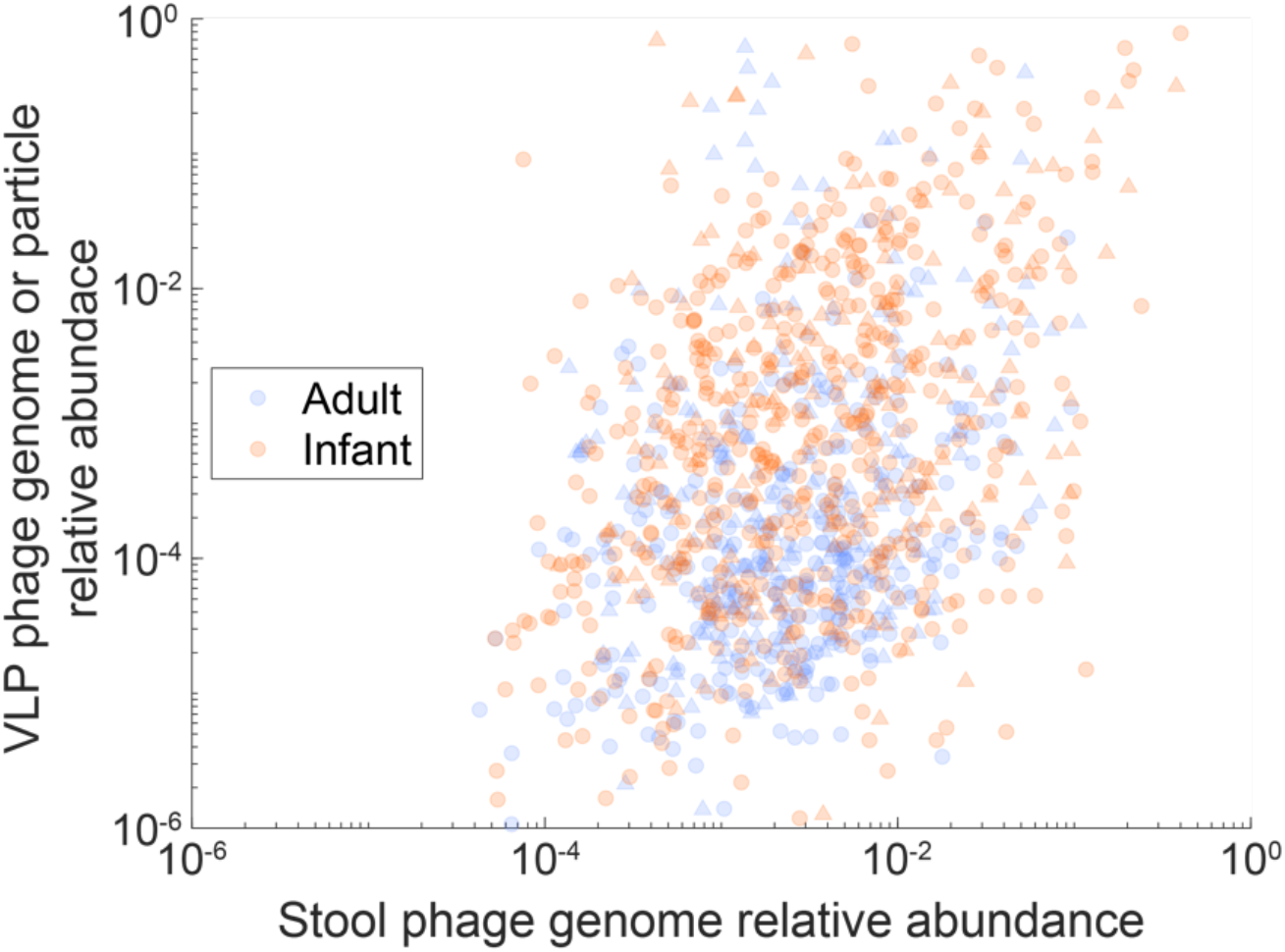
Species-level relative abundance analysis of the overlap between phage community quantifications using VLP- and stool-based approaches. Data and plotting methods are the same as **Fig. 1C**, except that relative abundance was used instead of absolute genome/particle density. This figure includes only phage shared between the VLP fraction and stool, corresponding to the central scatter plot of **Fig. 1C**. Relative abundance was defined relative to total taxonomic abundance of phage within VLPs/stool. Note that without the absolute abundance normalization employed in **Fig. 1C**, adult VLP abundances appear to be substantially lower than infant abundances.

**Figure S2:**
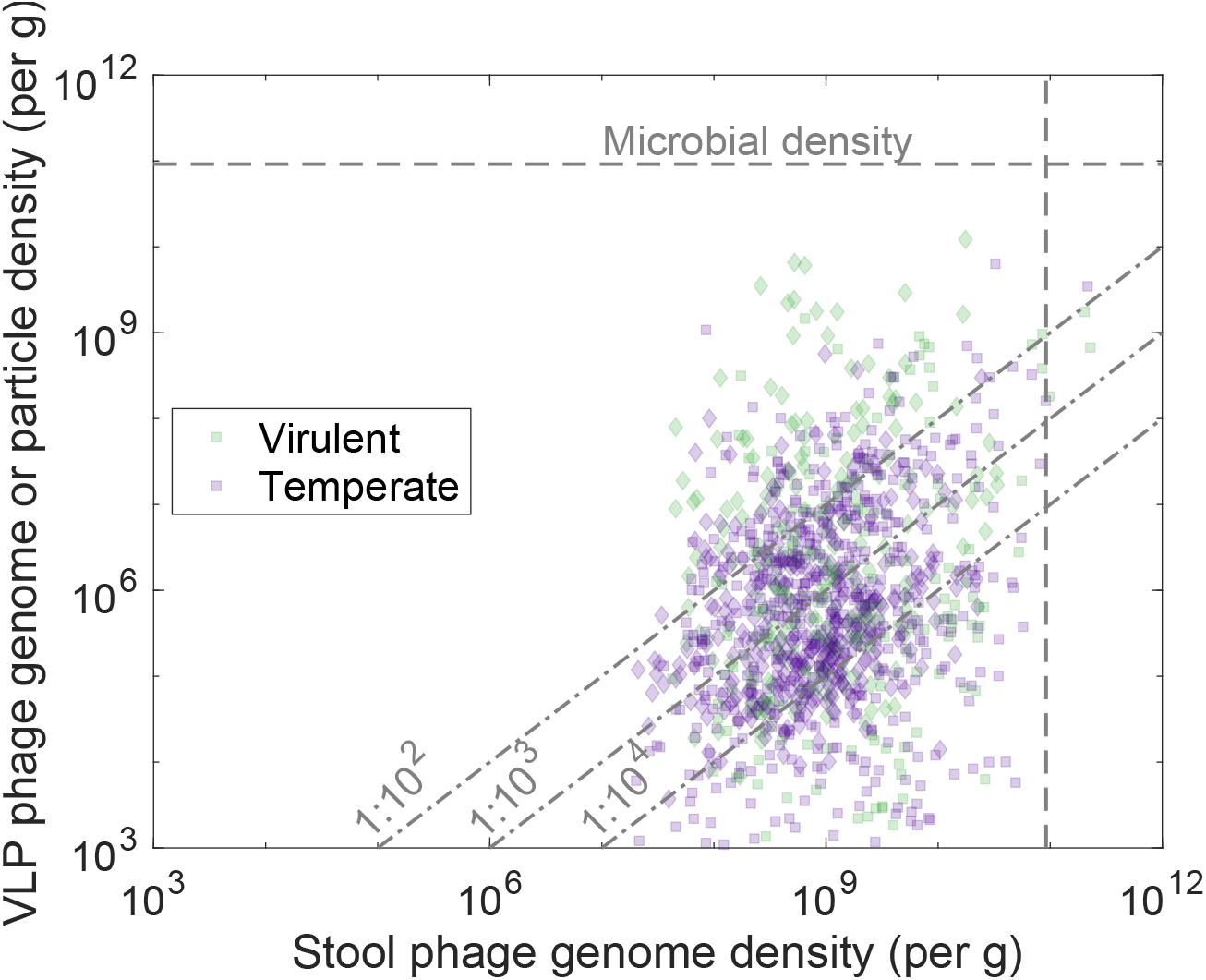
Species-level absolute abundance analysis of the overlap between VLP- and stool-based phage community quantifications, colored by predicted virulence. This figure is the same as **Fig. 1C**, except points are colored by predicted virulence rather than their dataset of origin. Diamond markers represent points from adult samples, while squares represent points from infant samples.

**Figure S3:**
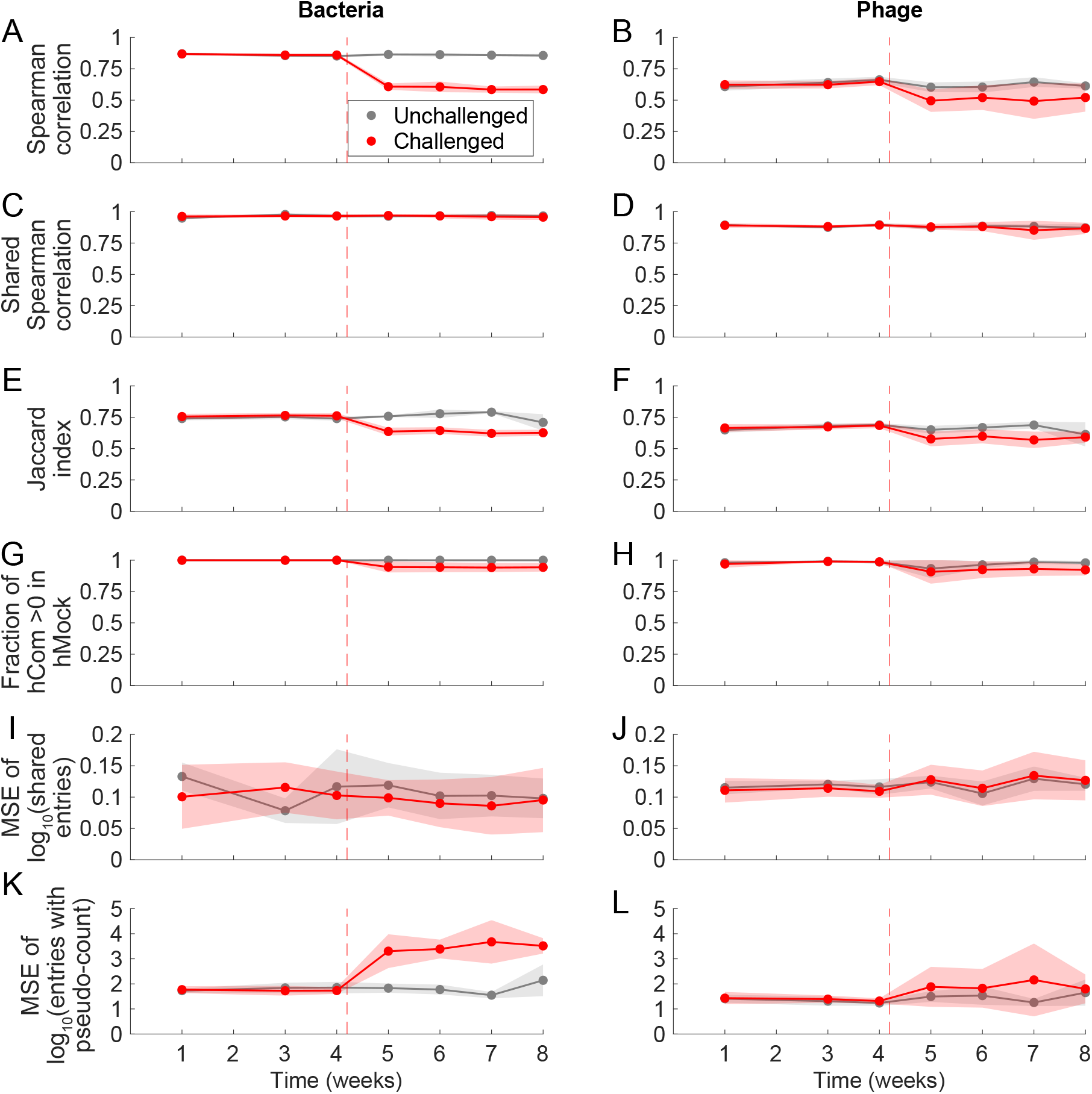
Time series of various reconstruction evaluation metrics in hCom2-colonized mice. Each row is a version of **Fig. 3C**,**D** but instead showing Spearman correlation of taxa read abundances (A,B), Spearman correlation of read abundances of species shared between hCom2/hMock sample pairs (C,D), Jaccard index (E,F), fraction of read abundances in hCom2-colonized mouse feces that are nonzero in hMock (G,H), mean squared error (MSE) of log_10_(read abundance) of species shared between hCom2/hMock sample pairs (I,J), and MSE of log_10_(read abundance) computed with a relative abundance pseudocount of 10^−7^ (K,L). All metrics were computed using species-level relative read abundances. Jaccard index is the number of shared species between an hCom2/hMock sample pair divided by the total number of species with nonzero abundance in at least one of the two samples. For (G,H), shared read abundances were normalized to the total bacterial or phage abundance in the hCom2-colonized mouse fecal sample.

**Figure S4:**
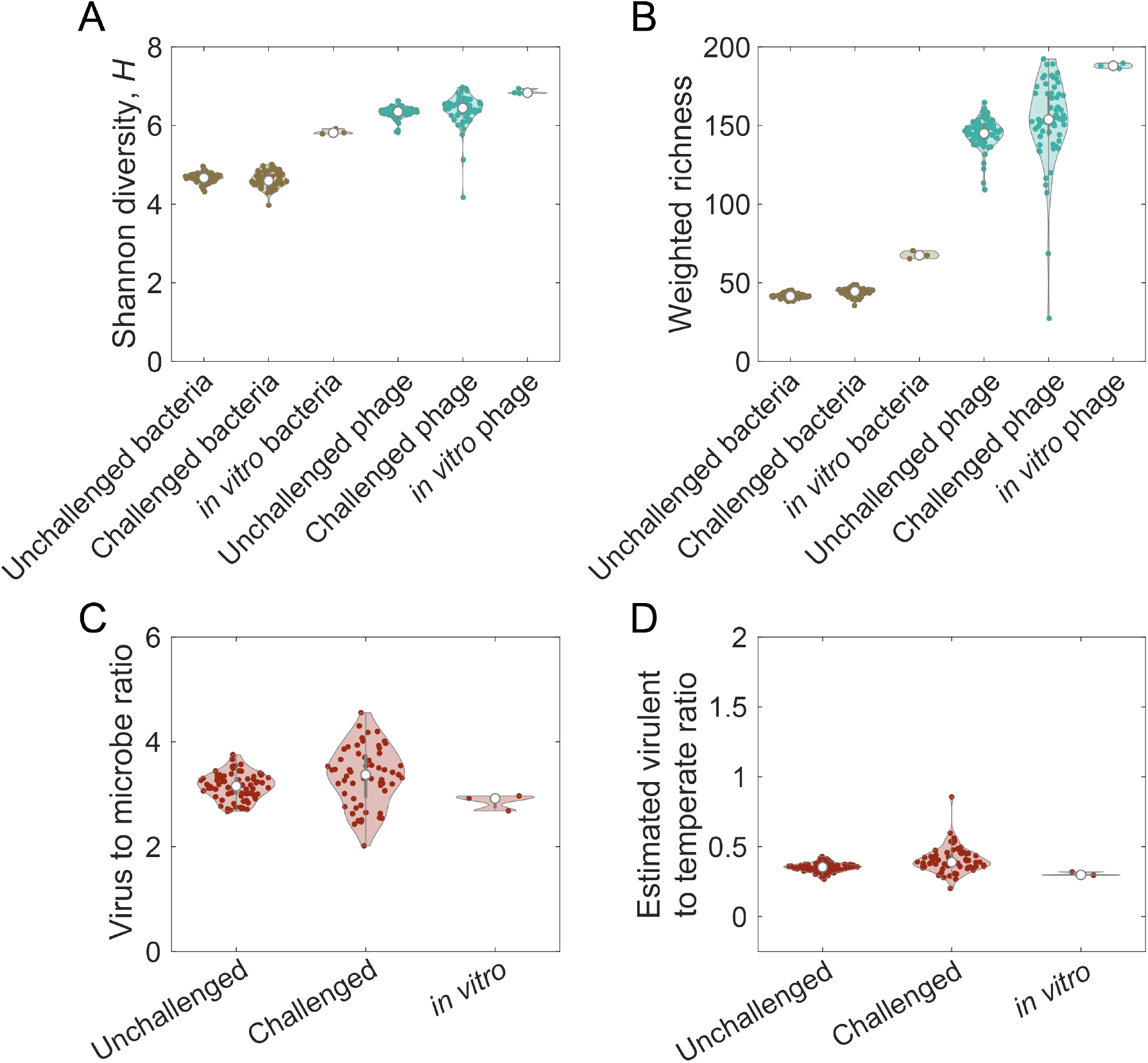
Comparison of phage and bacterial community properties in the feces of hCom2-colonized mice across conditions. Community properties and plotting methods are the same as in **Fig. 4**. ‘Unchallenged’ corresponds to *in vivo* samples from hCom2-colonized gnotobiotic mice that have not been exposed to a human stool sample, while ‘Challenged’ denotes samples from hCom2-colonized mice that have been exposed to a human stool challenge. ‘*in vitro*’ corresponds to hCom2 communities grown *in vitro*.

**Figure S5:**
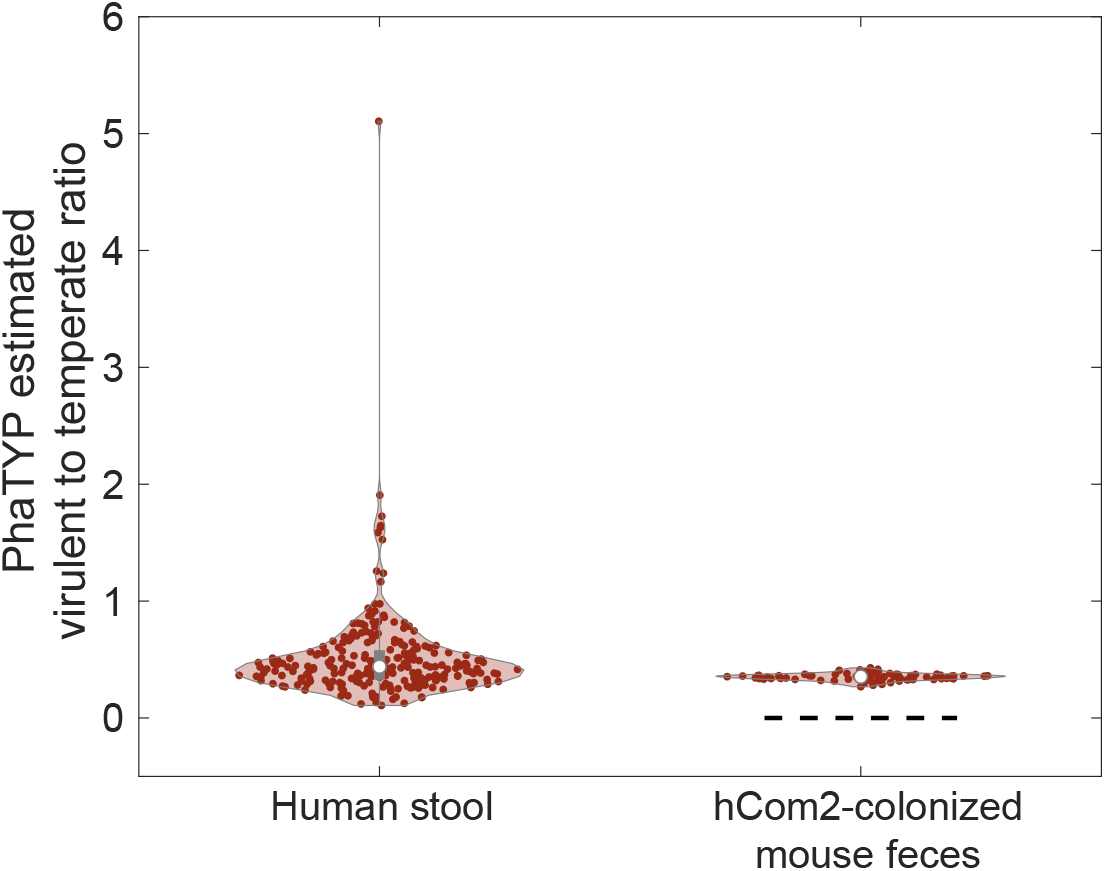
Comparison of the virulent-to-temperate ratio (VTR) between stool from humans and feces from hCom2-colonized mice, as estimated by PhaTYP. Equivalent to **Fig. 4D**, except the virulence prediction of phage genomes was performed using PhaTYP (58) instead of using the Phanta UHGV database (25). Dashed black line denotes the null expectation of VTR = 0 for a community constructed from axenic bacterial cultures.

